# LINE-1 transposon derived DNA lesions reshape the chromatin landscape to promote genome instability

**DOI:** 10.64898/2026.04.20.719662

**Authors:** Arun Kumar, Ankita Subhadarsani Parida, Kiran Kiran, Sunil K. Raghav, Bhavana Tiwari

**Affiliations:** Department of Biological Sciences, Indian Institute of Science Education and Research (IISER) Berhampur, Odisha, India; Immunogenomics & Systems Biology group, BRIC-Institute of Life Sciences (BRIC-ILS), Bhubaneswar, Odisha, India

**Keywords:** LINE-1 retrotransposon, ORF2p endonuclease, Double strand break, Genomic instability, Non homologous end joining (NHEJ), Enhancer remodelling

## Abstract

Aberrant expression of Long Interspersed Element-1 (LINE-1/L1) retrotransposons is increasingly linked to genomic instability in cancer, particularly in the context of compromised p53 function. The endonuclease (EN) domain of the LINE-1 ORF2 protein (ORF2p) generates DNA strand breaks, yet its role in shaping downstream chromatin and transcriptional responses remains poorly defined. Here, we investigate the impact of ORF2p-EN activity on DNA damage response signalling and epigenomic remodelling in A375 melanoma cells using a L1 expression system. ORF2p-EN led to activation of ATM dependent DNA damage signalling, evidenced by increased γH2AX accumulation. The EN induced DNA damage showed enrichment of yH2AX at fragile hotspot regions, consistent with localized DNA damage. We also observed increase in levels of the DNA repair factor XRCC5, suggesting a possible engagement of non-homologous end joining pathway. EN-induced DNA damage correlated with increased P300 activity and enhanced H3K27ac enrichment at regulatory regions, linking DNA damage signalling with chromatin acetylation. Pharmacological inhibition of ATM attenuated both γH2AX accumulation and H3K27ac levels, supporting a model in which chromatin remodelling occurring downstream of DNA damage signalling. Collectively, our findings position ORF2p endonuclease activity as an initiating source of DNA damage that is followed by chromatin remodelling and transcriptional reprogramming, with p53 acting as a critical modulator of this axis. These results provide mechanistic insight into how LINE-1 activation may contribute to oncogenic gene expression programs in cancer.

## Introduction

Long Interspersed Element 1 (LINE-1/ L1) retrotransposons are potent intrinsic sources of genomic instability (1). Although L1 is epigenetically silenced in most somatic tissues, it becomes atypically reactivated in several human cancers (2) where genome surveillance pathways are compromised (3,4). L1 elements occupy approximately 17% of the human genome (5). However, the majority are inactive, with only about 90-100 copies being retrotransposition competent where they retain the ability to replicate and insert themselves into new genomic locations (6). The full length L1 transcribes into a bicistronic mRNA that encodes two non-overlapping open reading frame (ORF) proteins, ORF1p and ORF2p. The 40 kDa ORF1p possess nucleic acid chaperonic activity (7). The 150 kDa ORF2p has two enzymatic activities of endonuclease (EN) and reverse transcriptase (RT) which are essential for retrotransposition (8). This process requires EN catalysed DNA nicking followed by reverse transcription of L1 mRNA into complementary DNA (cDNA), which is integrated into the genome (9–13). The DNA damaging potential of L1-ORF2p has been extensively characterized in the context of insertion mediated genomic rearrangements (14–16). The role of exogenous stressors such as radiation or chemotherapeutic agents in DNA damage responses is marked by γH2AX is well established (17). However, the consequences of DNA lesions generated by L1-ORF2p as an endogenous source of genome instability remains poorly understood. Studies have established that LINE-1 retrotransposition engages host DNA repair pathways. This process yields important insights into how DNA damage responses contribute to retrotransposition and insertion associated genome instability (18–22). Studies have linked replication stress (23) and transposon activation to chromatin remodelling (24), however, how ORF2p triggered DNA break is resolved by host repair is poorly understood. In light of DNA damage and enhancer regulation being linked processes (25), it is plausible that ORF2p induced DNA breaks occurring in transcriptional regulatory regions could contribute to chromatin remodelling in cancer genome (26).

L1 retrotransposition begins when EN nicks chromosomal target DNA (27,28) resulting in the accumulation of DNA damage marked by γH2AX foci, an early response to DNA strand breaks. H2AX is rapidly phosphorylated by ATM to decorate DNA break and spreads up to several mega bases from the original lesion site (29). Phosphorylation of H2AX is essential for recruitment of DNA repair proteins and promotes assembly of repair complexes at sites of damaged chromatin (30). However, if these double strand breaks (DSBs) are not resolved efficiently, it can cause chromosomal rearrangements and genomic instability, thereby promoting tumorigenesis (31–34). Activation of P300, a histone acetyltransferases leads to enhanced deposition of H3K27ac and chromatin remodelling (35) that further facilitates recruitment of DNA damage repair proteins (36) and transcription of genes such as MYC (37). Therefore, host DNA repair pathways are important for retrotransposition, but it is still unclear which specific repair pathways and if chromatin changes are triggered by ORF2p endonuclease induced DNA damage, especially when the DNA break does not result in successful insertion. Two major conserved pathways for DSB repair are non-homologous end joining (NHEJ) and homologous recombination (HR). NHEJ directly ligates broken DNA ends with limited sequence homology and is inherently error-prone, whereas HR uses the sister chromatid as a repair template during the S and G2 phases of the cell cycle. However, the choice between these pathways is strongly influenced by chromatin context, transcriptional status, and cell cycle stage (38–41). Several components of the NHEJ pathway, including XRCC4, XRCC5, and DNA-PKcs, have been implicated in facilitating transposon associated DNA repair and influencing retrotransposition efficiency (42,43). These findings indicate that host repair pathways, particularly NHEJ may modulate the efficiency and structural outcomes of L1 insertion events (44).

p53 mutant tumors can tolerate elevated levels of DNA breaks (45) and replication stress (46), thereby supporting tumor evolution by enabling adaptive transcriptional states. Importantly, p53 loss is known to derepress LINE-1 expression and impair DNA damage resolution (47). Given the critical interplay between transposable elements and host DNA repair machinery, it is essential to examine how retrotransposon induced DNA damage is repaired in cancer cells, particularly in p53^-/-^ backgrounds that are permissive to L1 derepression. In such contexts, defects in DNA damage sensing and repair pathway choice may create a permissive environ-ment in which ORF2p activity further destabilizes the genome. Although the individual roles of p53 loss and LINE-1 activation in genome instability are well established (48), the specific contribution of ORF2p endonuclease activity to DNA damage signalling, repair pathway engagement, and chromatin remodelling remains poorly defined.

In this study, we investigate how ORF2p endonuclease induced DNA damage intersects with chromatin remodelling and oncogenic transcriptional programs in melanoma cells. Using a LINE-1 expression system, we integrated RNA-seq with γH2AX ChIP-seq and H3K27ac ChIP-seq datasets to define endonuclease initiated DNA damage signalling. Our results link endonuclease triggered DNA damage signalling with transcriptional changes and followed by enhancer remodelling. We specifically demonstrate that ORF2p endonuclease activity induces DNA double strand breaks which trigger activation of ATM signalling, leading to γH2AX deposition at damaged chromatin. These damage associated regions correlates with *de novo* H3K27ac enrichment at regulatory elements. Notably, loss of p53 amplifies the magnitude and persistence of these chromatin changes. A substantial fraction of gained enhancers overlap with γH2AX marked loci, supporting a model in which ORF2p initiated DNA lesions possibly linked with enhancer remodelling and transcriptional reprogramming.

## Results

### L1-ORF2p endonuclease induces DNA damage in fragile hotspot sites

To investigate the impact of L1 ORF2p endonuclease (EN) on genomic stability we leveraged codon optimized full length human L1, pMT646 (L1 EN⁺) encoding both ORF1p and ORF2p. We also used pMT1093 (L1 EN⁻) with a point mutation of H230A in ORF2p rendering the EN domain catalytically inactive, as a control (49). A C-terminal 3×FLAG epitope tag was fused to ORF2p to facilitate its detection and immunoprecipitation. The constructs were expressed in wild-type (WT) or p53-deficient (p53^-/-^) A375 melanoma cells (50). Immunofluorescence was performed to validate the expression of ORF1p and ORF2p across both EN⁺ and EN⁻ expressing cells **(Figs. 1A and 1B, and S1A)**. Consistent with prior reports (51–53), ORF2p stained cells were observed in a subset of cells and interpreted qualitatively. Western blot was performed by probing γH2AX to measure the extent of DNA damage induced by EN domain of L1-ORF2p **(Figs. 1C and S1B)**. Quantification of γH2AX levels normalized to either β-actin **(Fig. 1D)** or to ORF2p **(Fig. S1C)**, revealed a marked increase in DNA damage in both WT and p53^⁻/⁻^ cells expressing L1 EN⁺ compared to L1 EN⁻, consistent with the role of p53 in DNA repair (54). Upon expression of EN domain of L1-ORF2p having a C terminal GFP fusion, we observed that it localized predominantly in the nucleus **(Fig. 1E)**. To assess whether this domain alone is sufficient to induce DNA damage, we performed comet assays on single cell nuclei transfected with EN-GFP and compared with empty vector controls. Cells expressing the endonuclease domain only, exhibited an increase in comet tail length, a direct measure of DNA fragmentation (**Figs. 1F and 1G**). These findings are in line with prior studies demonstrating that L1-ORF2p endonuclease domain introduces nicks in genomic DNA to initiate retrotransposition (55) which can be converted into double strand breaks during DNA replication or repair (56). Next, we performed ChIP-seq in WT and p53^⁻/⁻^ cells expressing either L1 EN⁻ or L1 EN⁺ using γH2AX. L1 EN⁺ expression induced elevated γH2AX deposition, yielding 4937 peaks in WT and 7453 in p53^⁻/⁻^ cells, whereas L1 EN⁻ expression generated only 1019 and 2109 peaks, respectively. These results demonstrate that ORF2p endonuclease induces DNA breaks, and this effect is amplified in the absence of p53, as shown by IGV snapshots at fragile sites FRA3B, FRA17A, FRAXC, and FRA22B **(Figs. 1H and S1D)**. Fragile sites are inherently prone to replication stress and breakage (57–60). These sites frequently overlap with large transcription units and regulatory regions, and DNA damage occurring at these loci can influence chromatin remodelling, and transcriptional regulation thereby linking genome instability to gene expression changes (61). Consistently, we observed elevated γH2AX enrichment at FRA3B-FCR and FRA3B-FDR in WT and p53^⁻/⁻^ A375 and U2OS (Osteosarcoma) cell lines expressing L1 EN⁺, whereas only modest or baseline levels were detected in the corresponding EN⁻ controls **(Figs. 1I and S1E)**. Notably, these fragile hotspot regions are typically AT-rich, which aligns with the known preference of the L1 endonuclease to introduce nicks at AT-rich target sequences (62,63). To further assess whether the observed enrichment of DNA damage at fragile sites is specific to L1 endonuclease activity, we compared yH2AX enrichment levels due to DNA damage induced either by cisplatin and endonuclease only domain. γH2AX ChIP-qPCR on cisplatin treated samples did not exhibit a similar locus-specific enrichment **(Fig. S1F),** suggesting that L1 endonuclease activity may display preferential targeting or accumulation of DNA damage at fragile genomic regions.

**Fig 1:**
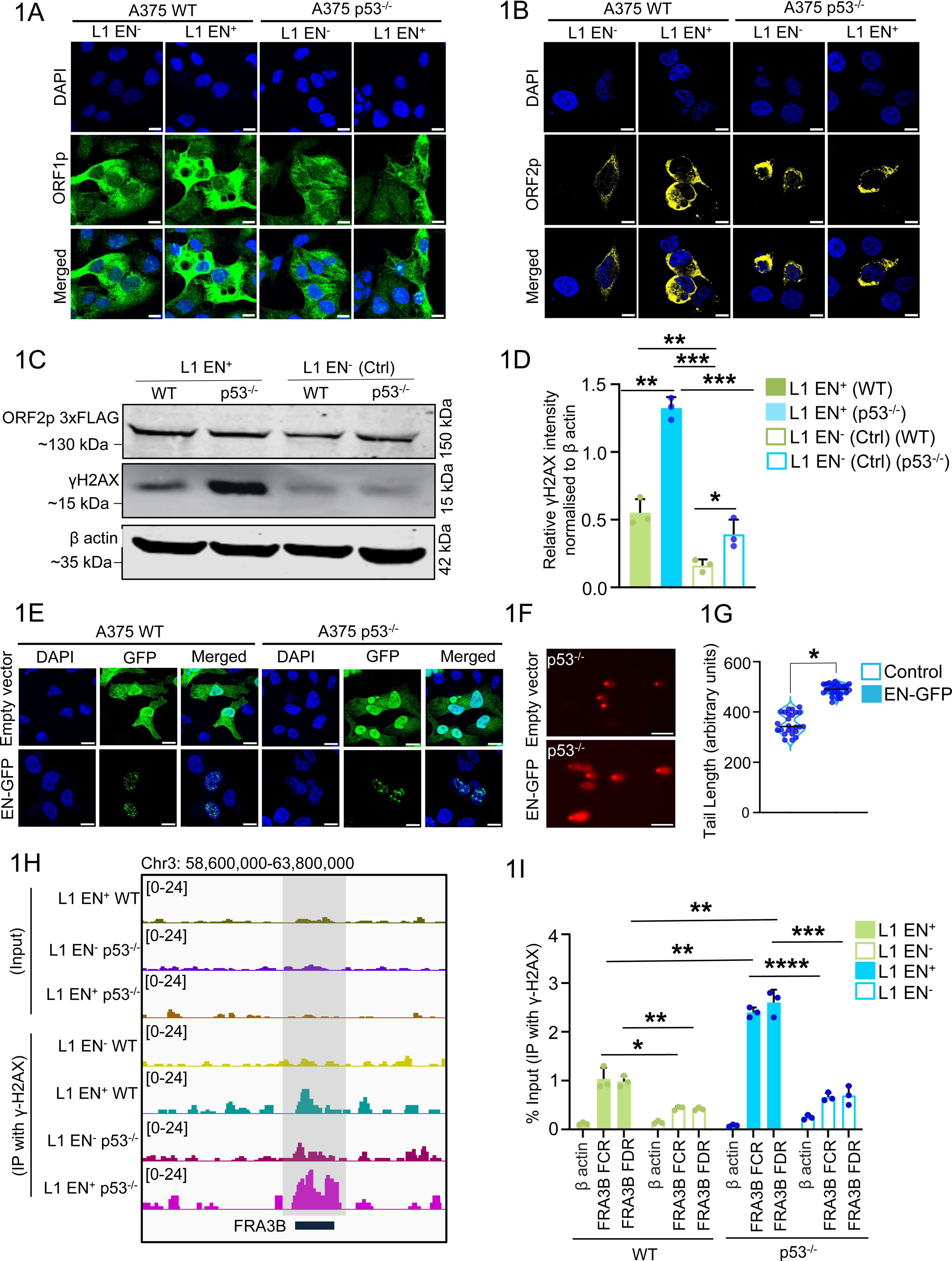
L1-ORF2p endonuclease domain induced DNA damage. Expression of codon optimized full length human L1-ORFeus constructs encoding ORF1 and ORF2. pMT646 encodes L1-ORF2p with a wild type endonuclease domain (L1 EN⁺), whereas pMT1093 encodes an ORF2p H230A endonuclease mutant (L1 EN⁻) carrying a point mutation that renders the endonuclease catalytically inactive. **(A-B)** Immunofluorescence analysis of L1 ORF1p and ORF2p in A375 wild type (WT) and p53 deficient (p53^-/-^) cell lines expressing pMT646 (L1 EN⁺) and pMT647 (L1 EN⁻) constructs, detected using (A) ORF1p (green) and (B) FLAG (yellow) tagged antibodies and nucleus counterstained with DAPI (blue). Scale bar represents 5µm. **(C)** Western blot demonstrates expression of ORF2p from both L1 EN⁺ and L1 EN⁻ (Ctrl) constructs in A375 WT and p53^⁻/⁻^ cells using anti FLAG antibody. The γH2AX bands measure the endonuclease domain triggered DNA damage across the samples. β actin served as a loading control. **(D)** Densitometric quantification of γH2AX band intensities from immunoblots in A375 WT and p53⁻/⁻ cells expressing L1 EN⁺ and L1 EN⁻ constructs. The normalization of γH2AX is performed with both L1 ORF2p as well as β-actin bands. The Fiji quantified data represent mean ±S.D. across three independent biological replicates. Statistical significance was evaluated using the Mann Whitney U test, with significance levels represented as follows: p*<0.05, p**<0.01 and ns = non-significant. **(E)** Immunofluorescence images of A375 WT and p53^⁻/⁻^ cells expressing either empty vector or a C terminal GFP tagged endonuclease (EN) domain of L1 ORFeus ORF2p (CMV::EN-GFP). EN-GFP is driven by CMV promoter (green). Nuclei are counterstained with DAPI (blue). Scale bar, 5 µm. **(F, G)** Representative images from fluorescence microscopy (**F**) and quantification of tail length (**G**) from A375 p53^-/-^ cells transfected with either empty vector (EV) or CMV::EN-GFP. DNA damage was assessed by alkaline comet assay and analyzed using Fiji software (n = 50 cells in each condition, 3 independent experiments). Data are presented as mean ± SEM. Statistical significance was determined using an unpaired two tailed Student’s t test; p < 0.05. Scale bar represents 5µm. **(H)** IGV snapshots showing γH2AX ChIP-seq signal at the common fragile site FRA3B across L1 EN⁻ and L1 EN⁺ conditions in WT and p53^⁻/⁻^ A375 cells. Tracks represent normalized γH2AX ChIP-seq coverage (RPM). Shaded regions indicate the FRA3B locus. **(I)** Chromatin immunoprecipitation (ChIP) followed by qPCR was performed to assess the enrichment of γH2AX at fragile sites FRA3B of FCR and FDR region in A375 cells transfected with either L1 EN⁺ or L1 EN⁻. β-actin served as a negative control. Data are presented as mean ± S.D across three independent biological replicates. Statistical analysis was conducted using the Mann Whitney U test; significance is indicated as p* < 0.05, p**<0.01, p***<0.001 and ns= non-significant.

### LINE-1 endonuclease mediated DNA damage reshapes transcriptional programs

To assess the change in transcriptional program due to expression of L1-ORF2p endonuclease we performed RNA-seq on A375 WT and p53^⁻/⁻^ melanoma cells expressing either L1 EN⁺ or EN⁻. Principal component analysis (PCA) of three biological replicate samples revealed tight clustering conditions in WT and p53^-/-^ cell lines expressing L1 EN^-^ and L1 EN^+^. One outlier from p53^⁻/⁻^ EN⁺ group was excluded from downstream analyses to preserve data integrity (**Fig. S2A**). Differential gene expression analyses **(Figs. 2A-C)** revealed significant and genotype specific transcriptional alterations upon L1 EN⁺ expression relative to L1 EN⁻ across A375 WT and p53^⁻/⁻^ cells. In WT cells **(Fig. 2A)**, 560 genes were significantly upregulated including DNA repair genes such as FANCD2, MSH2, and XRCC5, while 197 genes were downregulated. In p53^-/-^ **(Fig. 2B)**, 193 genes were significantly upregulated including MYC, FANCD2, and XRCC5, while 198 genes were downregulated. The comparative volcano plot revealed increased expression of components associated with the NHEJ pathway, an error-prone DNA repair pathway and oncogenic programs in both p53^⁻/⁻^ EN⁺ and WT EN⁺ conditions. Differential gene expression analysis visualized in the heatmap showed distinct transcriptional differences between EN⁺ and EN⁻ expressing cells in both genetic backgrounds. Notably, EN⁺ expression led to higher transcript levels of genes encoding components of NHEJ pathway, including XRCC4, XRCC5, and NHEJ1, alongside increased expression of genes associated with alternative repair and replication stress responses, such as NBN, FANCD2 **(Fig. 2C, Supporting Table 2)**. RT qPCR validation demonstrated modest upregulation of the NHEJ factors XRCC5 and LIG4, whereas only minor changes were observed for XRCC2 and MRE11, and no significant alteration was detected in RAD51 expression. These results indicate selective transcriptional remodelling of genes involved in double-strand break repair in response to L1 EN⁺ expression in WT and p53^-/-^ **(Figs. S2B and S2C).** These changes are observed due to expression of EN driven by either native endogenous L1 5′UTR promoter or CMV promoter **(Figs. 2D and S2E)**. This demonstrates that the effect is attributable to ORF2p EN domain mediated DNA damage rather than to retrotransposition. These results also show that expression of EN led to elevated levels of XRCC5 and LIG4 transcripts in both A375 WT **(Fig. 2D)** and p53^⁻/⁻^ **(Fig. S2E)** cells, supporting a direct role for EN in modulating DNA repair pathway choice. We observed elevated XRCC5 levels in both WT and p53^-/-^, consistent with persistence of DNA lesions and enhanced transcripts of NHEJ pathway genes **(Fig. S2D)**. However, we did not observe the similar transcriptional changes when we expressed APE1 endonuclease, indicating that L1-ORF2p endonuclease elicits a unique transcriptional programs of DNA damage repair rather than a generic endonuclease driven effect **(Fig. 2E)** (64). This suggests that DNA breaks generated by L1-ORF2p endonuclease elicit a transcriptional program for NHEJ associated components. Gene set enrichment analysis (GSEA) in both A375 WT and p53^⁻/⁻^ cells expressing L1 EN⁺ revealed strong enrichment of gene sets associated with DNA damage response, DNA metabolic processes, double strand break repair, and NF-κB signalling across both genotypes **(Figs. 2F-I, S2F-I)**. Moreover, pathway enrichment analysis comparing L1 EN⁻ and L1 EN⁺ in p53^⁻/⁻^ cells confirmed that ORF2p-EN functions as a potent inducer of genotoxic stress **(Fig. S2J)**. Collectively, these findings suggest that L1-ORF2p endonuclease activity elicits a coordinated DNA damage associated transcriptional program, since we observed heightened effects in p53^-/-^ as compared to the WT, we think that loss of p53 amplifies deregulation of repair related gene expression.

**Fig 2:**
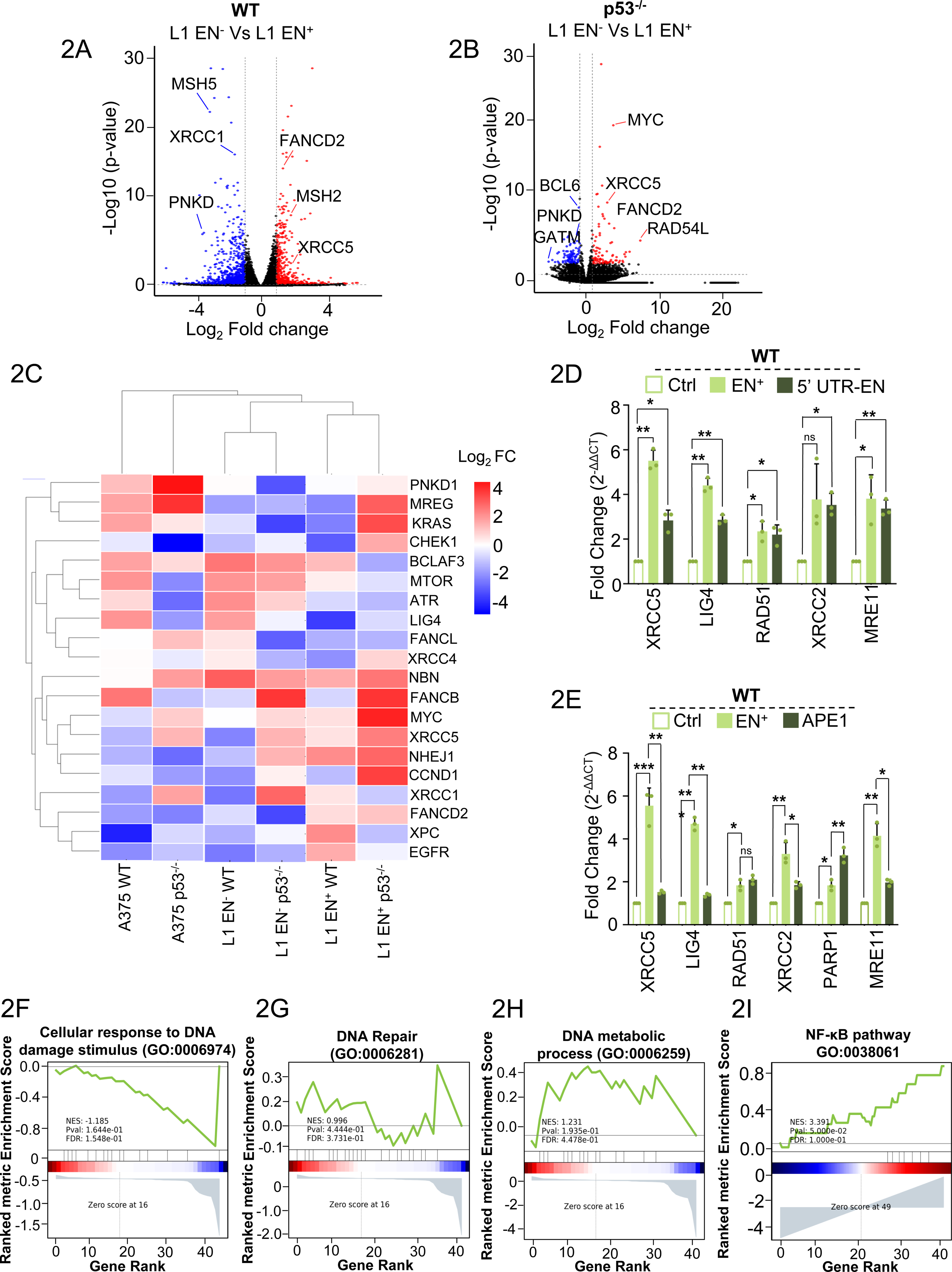
Transcriptomic analysis in A375 WT and p53^-/-^ cells expressing L1 EN^+^ and L1 EN^-^. **(A,B)** Volcano plots depicting differential gene expression in A375 melanoma cells expressing different L1 constructs. (A) WT A375 cells expressing L1 EN⁻ versus EN⁺ constructs. (B) p53^⁻/⁻^ A375 cells expressing L1 EN⁻ versus L1 EN⁺ constructs. The x-axis represents log₂ fold change in gene expression (EN⁺ relative to EN⁻), and the y-axis shows -log₁₀(p value) derived from differential expression analysis. Genes are color coded based on statistical significance and direction of change: upregulated genes (red), downregulated genes (blue), and non-significant genes (black). **(C)** Heatmap illustrating the expression patterns of 20 selected genes across six experimental conditions in A375 lines, untransfected WT, p53^⁻/⁻^ and WT, p53^⁻/⁻^ transfected either with L1 EN^+^ or L1 EN^-^ (Ctrl), each represented by three biological replicates. Gene expression values are shown as scaled expression levels, encompassing both positive and negative values, and visualized using a diverging color scale centered at zero (red indicates high expression, blue indicates low or negative expression, and white represents baseline expression). Hierarchical clustering of both genes (rows) and samples (columns) was performed to identify shared expression patterns and to emphasize similarities and differences in transcriptional responses across L1 endonuclease activity states and p53 genetic backgrounds. **(D)** Bar plots displaying average mRNA expression levels using qRT PCR (mean ± S.D.) for XRCC5, LIG4, RAD51, XRCC2, and MRE11 across three independent biological replicates in WT A375 cell lines expressing Empty-vector (control), CMV-EN-GFP, endonuclease domain c-terminally fused with GFP is expressed under native promoter of L1 (5’UTR-EN-GFP). Statistical significance was assessed using the Mann Whitney U test. Significance levels are indicated as follows: *p < 0.05, **p < 0.01, ***p < 0.001, and ns = not significant. **(E)** Bar plots displaying average mRNA expression levels by qRT-PCR for for XRCC5, LIG4, RAD51, XRCC2, PARP1 and MRE11 in WT A375 cell lines expressing empty vector control, APE1 expressed under CMV promoter fused c-terminally with GFP and CMV-EN-GFP. The individual data points indicate three biological replicates, mean ± S.D., statistical significance was assessed using the Mann Whitney U test. Significance levels are indicated as follows: *p < 0.05, **p < 0.01, ***p < 0.001, and ns = not significant. **(F-I)** Gene Set Enrichment Analysis (GSEA) plots comparing A375 p53^⁻/⁻^ cells expressing L1 EN⁻ versus L1 EN⁺ constructs. **(F)** GO:0006974 - Cellular response to DNA damage stimulus, showing enrichment in L1 EN⁻ relative to L1 EN⁺ conditions (NES = -1.251, FDR = 5.80 × 10⁻²). **(G)** GO:0006259 - DNA metabolic process, displaying modest enrichment between L1 EN⁻ and L1 EN⁺ conditions (NES = 1.034, FDR = 7.73 × 10⁻¹). **(H)** GO:0006281 - DNA repair, showing significant enrichment in the L1 EN⁻ background, with gene set members indicated by black vertical bars along the ranked gene list and the distribution of enrichment signals shown in the lower heatmap. **(I)** GO:0038061 - NF-κB signaling pathway, enriched in L1 EN⁻ versus L1 EN⁺ p53^⁻/⁻^ cells (NES = -1.390, FDR = 4.00 × 10⁻²). In each plot, the green curve represents the running enrichment score, reflecting the degree to which genes in the indicated pathway are overrepresented at the extremes of the ranked gene list derived from differential expression analysis.

### L1-ORFeus2p expression alters native L1 transposon landscape and genome stability features

Building upon the transcriptional remodelling associated with codon optimized (L1-ORFeus) L1-ORF2p endonuclease activity, we next evaluated the impact of L1-ORFeus expression on native retrotransposon expression and activity. RNA sequencing of A375 WT and p53^-/-^ cells expressing either L1 EN⁺ or EN⁻ ORF2p revealed increased expression of native retrotransposon families, including LINE-1, LTRs and Alu elements, in the presence of L1 EN⁺ regardless of p53 status. This is illustrated by violin plots comparing EN⁺ and EN⁻ conditions in both WT **(Fig. 3A)** and p53^-/-^ **(Fig. 3B)** A375 cells (**Fig. S3A),** indicating that endonuclease activity of synthetic L1-ORFeus ORF2p could enhance native retrotransposon transcript levels. This effect was particularly pronounced for L1 subfamilies, supporting the notion that the trans effect of ORFeus retrotransposition machinery can promote native L1 expression levels (65). The increased levels of Alu elements further supports that it can exploit the L1 retrotransposition machinery for their own propagation (66). A heatmap of differentially expressed retroelement subfamilies highlighted robust transcriptional enrichment of L1s, including L1Hs, L1PA2, L1PA3 and LTRs specifically in L1 EN⁺ expressing cells (**Fig. 3C, Supporting Table 3**). Strikingly, this activation was markedly intensified upon p53 loss (67,68), in agreement with earlier reports that p53 constrains retroelements through epigenetic repression.

**Figure 3:**
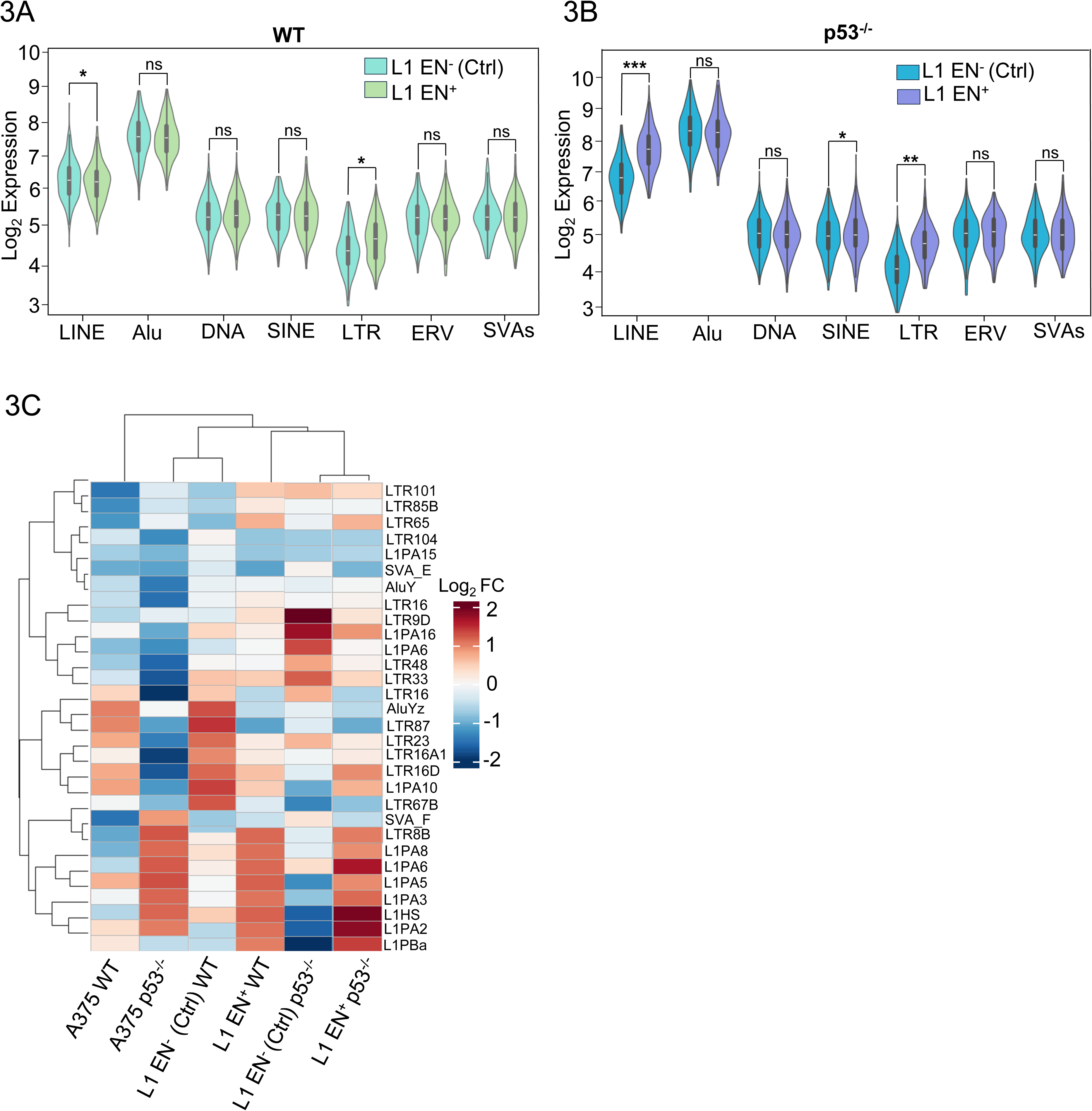
Transcriptomic analysis of native transposons in L1 EN^-^ vs L1 EN^+^. **(A,B)** Violin plots showing the expression distribution of selected transposable element (TE) families in A375 WT cells (A) and p53⁻/⁻ cells (B), comparing L1 EN⁻ and L1 EN⁺ expression. Each violin represents data from three biological replicates, with individual dots indicating replicate values and the embedded box plot marking the median and interquartile range. Statistical significance was evaluated using Fisher’s exact test, with “ns” indicating non-significant differences (p > 0.05), *p<0.05. **(C)** Heatmap showing top 30 differential expression of individual TE loci across all experimental conditions. Rows represent TE loci annotated by subfamily and each column represented by three biological replicates of A375 cell line samples (WT, p53^-/-^, L1 EN^-^ WT, L1 EN^+^ WT, L1 EN^-^ p53^-/-^, L1EN^+^ p53^-/-^). The colour gradient reflects relative expression levels, upregulated (red) to downregulated (blue). Hierarchical clustering (Euclidean distance, complete linkage).

### LINE-1 ORF2p induced DNA damage signalling precedes epigenetic change

As described in the previous result **(Fig. 1)**, expression of L1 EN⁺ resulted in increased DNA damage, evidenced by elevated γH2AX accumulation. We next performed co-localization analysis of EN-GFP and γH2AX. Co-localization between EN and γH2AX foci was observed in both WT and p53^⁻/⁻^ cells, however, the degree of spatial overlap was reduced in p53^⁻/⁻^ cells despite exhibiting a higher number of γH2AX foci **(Fig. 4A)**. This difference in p53^-/-^ could be due to potential influence of p53 loss on cell cycle distribution and its impact on DNA repair that requires further investigation.

**Result 4:**
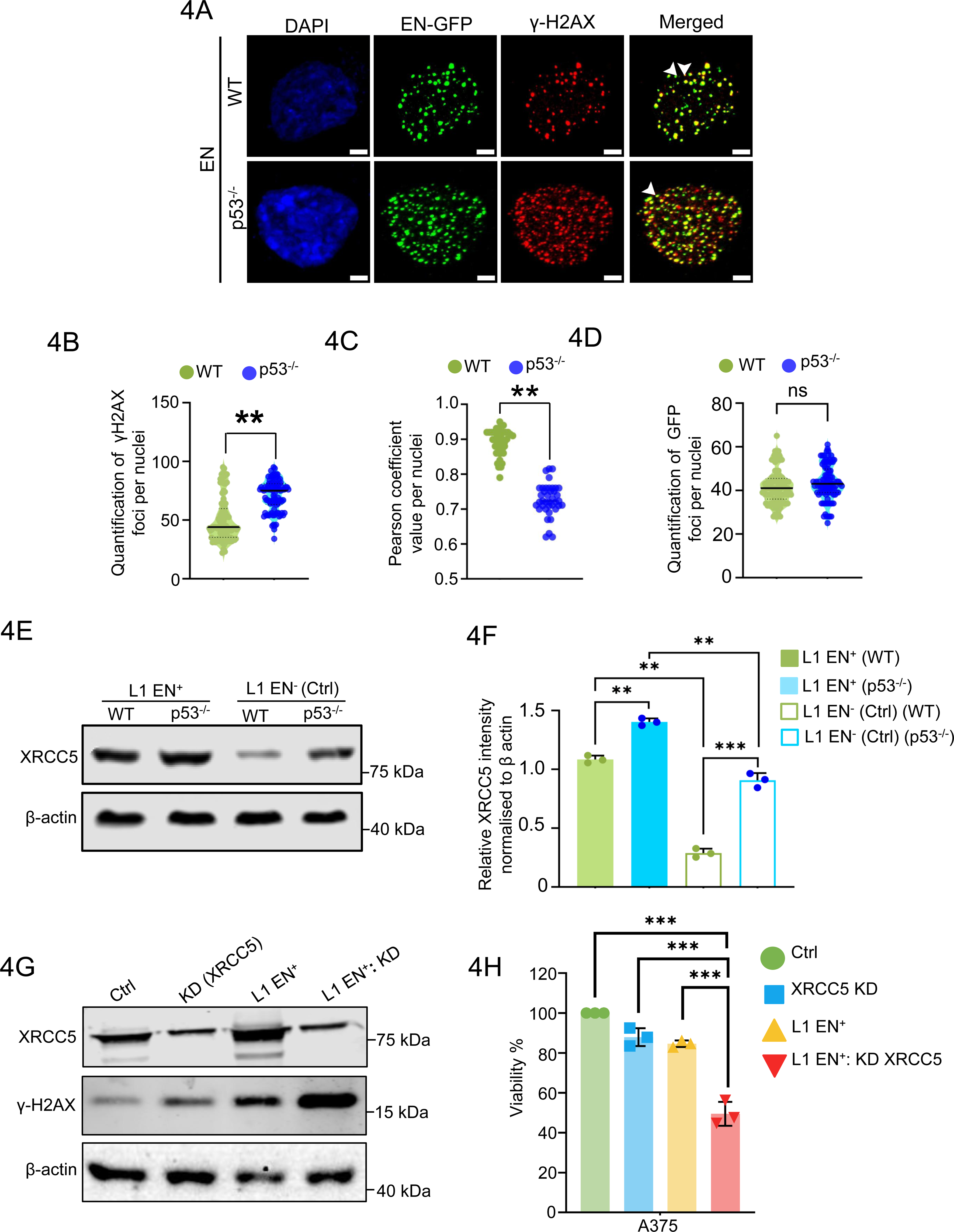
Analyzing DNA damage in EN-GF P expressed cells. **(A)** Representative confocal images of A375 WT and p53^⁻/⁻^ cells transfected with CMV-EN-GFP (EN-GFP, green), immunostained for the DNA damage marker γH2AX (red), and counterstained with DAPI (blue) to visualize nuclei (A). Immunofluorescence was performed to assess spatial co-localization of EN-GFP and DNA damage foci. EN was detected using an anti-GFP antibody, while γH2AX was visualized using an anti phospho-H2AX (Ser139) antibody. Scale bar: 2 μm. **(B)** Quantitative analysis of γH2AX foci in CMV-EN-GFP positive WT and p53^-/-^ A375 cells. For each condition, a total of 50 GFP positive nuclei (∼5 nuclei per field across 10 randomly selected fields per replicate) were analyzed from three independent biological replicates. Only γH2AX foci with diameters between 1.0 and 1.5 μm were included in the analysis. Statistical significance was assessed using the Mann Whitney U test, with **p < 0.01 considered significant, ns = not significant. **(C)** Dot plot representing the co-localization analyses of CMV-EN-GFP and γH2AX in A375 WT and p53^⁻/⁻^ cells. The Pearson correlation coefficients (r) range from 0.85 to 0.95 in A375 WT cells and from 0.65 to 0.85 in A375 p53^⁻/⁻^ cells. Statistical significance was assessed using the Mann Whitney U test, with **p < 0.01. **(D)** Violin plot overlaid with individual data points showing the number of GFP puncta in GFP positive nuclei. A total of 100-50 GFP positive nuclei (∼5 nuclei per field across 30 fields per replicate) from three independent biological replicates were analyzed. Statistical significance was assessed using the Mann Whitney U test. ns = non-significant. **(E)** Western blot of XRCC5 protein expression in A375 WT and p53^-/-^ cells L1 EN⁺ and L1 EN⁻. β-actin served as the loading control. Control is abbreviated as Ctrl. **(F)** Densitometric quantification of XRCC5 protein levels from Western blots in A375 WT and p53^-/-^ cells expressing L1 EN⁺ and L1 EN⁻. Expression levels were normalized to β-actin, and individual data points indicate each biological replicate where mean ± SEM across three biological replicates. Statistical significance was evaluated using the Mann Whitney U test, with significance levels represented as follows: *p<0.05, and ns = non-significant. **(G)** Representative immunoblot analysis of A375 cells transiently transfected with pTLVC2 vector control, pTLVC2-CRISPR targeting XRCC5, L1 EN^+^, or co-transfected with XRCC5 targeting CRISPR and EN^+.^ Whole cell lysates collected 48h post transfection were probed for XRCC5 to assess depletion efficiency and for γH2AX as a marker of DNA double strand breaks. β-actin was used as a loading control. **(H)** MTT assay showing cell viability of A375 cells transfected with either pTLVC2 control, or pTLVC2-CRISPR sgRNA targeting XRCC5. A375 cells were co-transfected with both pTLVC2 carrying sgRNA and EN^+.^ The combined perturbation results in a marked reduction in viability compared with either condition alone. Data are presented as mean ± SD from three independent experiments. Statistical significance was assessed using appropriate two-tailed tests.

Next, to determine whether reduced co-localization reflected differences in EN abundance instead of altered damage response, intensities of nuclear EN-GFP and γH2AX foci were quantified per cell basis, and co-localization was evaluated within matched EN intensity bins across genotypes **(Fig. 4B-D)**. Across equivalent EN expression ranges, p53^⁻/⁻^ cells consistently exhibited reduced spatial coincidence between EN-positive foci and γH2AX, indicating that the observed phenotype was not attributable to EN overabundance or signal saturation. Despite robust γH2AX accumulation in p53^⁻/⁻^ cells expressing EN domain, the spatial coincidence between EN positive foci and γH2AX foci was significantly reduced compared to WT cells. Similar observations were seen when EN-GFP was expressed under native L1-5UTR promoter **(Fig. S4A)**.

L1 EN⁺ generates DSBs that are presumed to engage canonical DNA repair pathways; however, the extent to which these lesions are accurately sensed and processed remains unclear. NHEJ is initiated by recruitment of XRCC5 heterodimer to DNA ends (69). Western blotting experiments demonstrated a modest increase in the level of XRCC5 (**Fig. 4E and F**), a component of NHEJ pathway in both WT and p53^⁻/⁻^ A375 cells expressing L1 EN^+^ as compared to EN^-^ controls. Next we visualised the co-localisation of either XRCC5 or RAD51 with γH2AX in A375 WT and p53^⁻/⁻^ cells expressing either L1 EN⁺ and L1 EN⁻. In L1 EN⁺ expressing cells, γH2AX foci co-localized with XRCC5 in both WT and p53^⁻/⁻^ conditions; however, this colocalization was significantly enhanced in p53^⁻/⁻^ cells compared to WT. In contrast, colocalization of γH2AX with RAD51, a key component of homologous recombination (HR) mediated repair (68), remained minimal in WT A375 cells expressing L1 EN⁺ **(Fig. S4B-E).** Here we note that XRCC5-γH2AX spatial proximity is a qualitative and supportive evidence of recruitment of repair associated factors to sites of DNA damage, rather than as a mechanistic readout of NHEJ activity. Additionally, the difference in p53 deficient could be due to difference in altered canonical pathways governed by p53 including cell cycle arrest and DNA damage. To functionally examine the role of the NHEJ factor XRCC5 (Ku80) in the context of EN-induced DNA damage, we performed CRISPR-Cas9 mediated targeting of XRCC5 in A375 cells expressing L1 EN⁺. Reduced XRCC5 levels were confirmed by immunoblotting, indicating efficient but incomplete disruption (**Fig. S4F**). EN expression alone led to increased γH2AX levels, consistent with induction of DNA double strand breaks. Notably, partial loss of XRCC5 in EN expressing cells led to increase in the levels of γH2AX, indicating impaired repair of DNA lesions in these cells (**Fig. 4G**). We next assessed the consequences of XRCC5 perturbation on cell viability using an MTT assay (**Fig. 4H**). XRCC5 targeting alone led to a modest reduction in viability, whereas combined EN expression and XRCC5 targeting resulted in an approximately 50% decrease in cell viability, demonstrating that XRCC5 mediated repair is required for cellular tolerance to LINE-1 induced genomic stress.

To further investigate the DNA damage response following EN induced DNA damage, we assessed activation of DNA repair proteins and chromatin associated marker at different time points. A schematic overview **(Fig. 5A)** outlines the molecular events following EN mediated DNA cleavage. EN expression led to rapid accumulation of γH2AX, followed by increased activation of 53BP1 and XRCC5. Following these events, we also observed increase in levels of H3K27ac, a marker of transcriptionally active chromatin, suggesting a link between EN triggered DNA damage to changes in chromatin. Western blot analysis confirmed a time dependent increase in γH2AX, 53BP1, XRCC5, and H3K27ac at 24 and 48 hours post transfection relative to baseline **(Fig. 5B and C)**. To understand mechanistic basis of XRCC5 activation, we analysed accumulation of XRCC5 in the chromatin enriched fraction due to EN induced DNA damage. A schematic model **(Fig. 5D)** proposes that XRCC5 transitions from a soluble nuclear pool to a chromatin bound state upon DNA damage. Fractionation based Western blot analysis **(Fig. 5E and F)** confirmed significant enrichment of XRCC5 in the chromatin fraction following EN expression, indicating its recruitment to sites of DNA damage. In parallel, increased levels of CBP/P300 as a transcriptional co-activator and chromatin regulator (70) and elevated H3K27ac levels were observed in chromatin fractions suggesting that EN induced DNA damage promotes histone acetylation and enhancer activation. These changes are likely reinforced by transcriptional upregulation of factors such as MYC and NF-κB, linking DNA damage signalling to transcriptional and epigenetic reprogramming as observed in **Fig 2**.

**Result 5:**
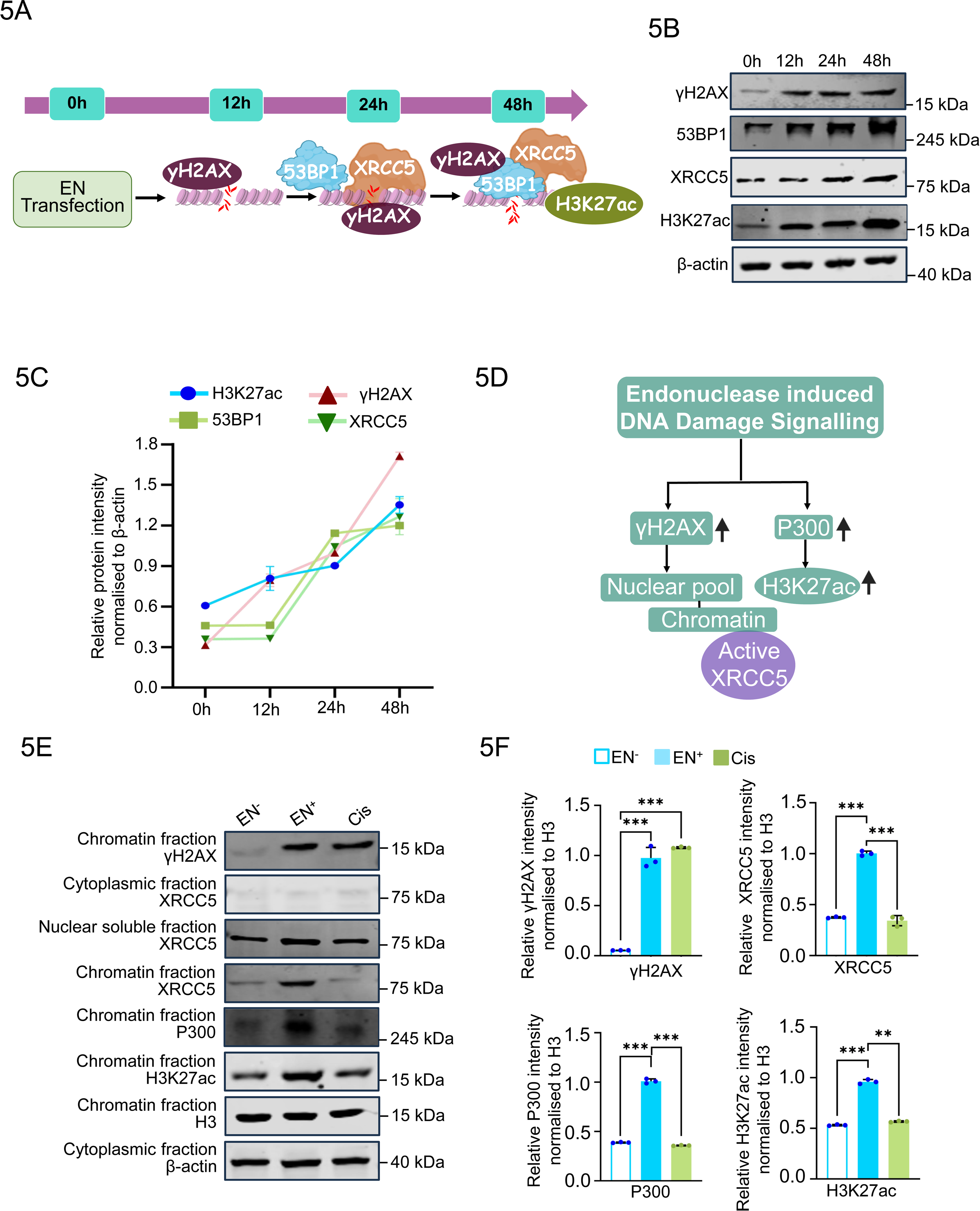
DNA damage Signalling response to LINE-1 endonuclease (EN) expression. **(A)** Schematic overview of temporal activation of DNA damage signalling factors triggered by LINE-1 endonuclease (EN) expression. EN-induced DNA damage is marked by γH2AX formation, accompanied by recruitment of DNA repair factors 53BP1 and XRCC5, followed by increase in H3K27ac levels. **(B)** Western blot analysis of γH2AX, 53BP1, XRCC5, and H3K27ac expression at 0, 12, 24, and 48 hours following EN expression in A375 WT cells. **(C)** Quantitative densitometry analysis of γH2AX, 53BP1, XRCC5, and H3K27ac protein levels over time (0, 12, 24, and 48 hours) after EN expression. Protein levels were normalized to β-actin, with individual data points representing biological replicates. Data are presented as mean ± SD from three independent experiments. **(D)** Schematic representation of the redistribution of γH2AX, XRCC5, P300, and H3K27ac from the nuclear soluble fraction to chromatin in response to EN induced DNA damage. **(E-F)** Western blot analysis of cytoplasmic, nuclear soluble, and chromatin fractions to determine the distribution of γH2AX, XRCC5, P300, and H3K27ac. Histone H3 and β-actin were used as chromatin and cytoplasmic loading controls, respectively. Band intensities were quantified by densitometry, and statistical significance was assessed using the Mann-Whitney U test (*p < 0.05; ns, not significant). Data are presented as mean ± SD from three independent biological replicates, with individual data points shown.

### Chromatin remodelling induced by L1-ORF2p endonuclease activity

DSBs induced by genotoxic stress are known to initiate a broad chromatin response, often marked by γH2AX spreading across megabase scale domains, which can influence the activity of nearby regulatory elements and alter gene expression programs (71–74). Given that L1-ORF2p endonuclease activity generates DSBs, we next sought to examine histone modification patterns associated with active chromatin. Western blot analysis revealed an increase in H3K27ac levels in WT and p53^⁻/⁻^ cells expressing L1 EN⁺ construct as compared to EN⁻ counterparts **(Fig. 6A and 6B)**. This increase was consistently observed across biological replicates indicating that ORF2p endonuclease activity might be associated with enhanced histone acetylation at active chromatin regions. To further confirm if underlying changes in chromatin is triggered due to EN, we performed chromatin immunoprecipitation followed by sequencing (ChIP-seq) using antibodies against H3K27ac and H3K4me3 in A375 WT and p53^⁻/⁻^ cells expressing L1 EN⁺, along with p53^⁻/⁻^ cells expressing L1 EN⁻ **(Figs. 6-8 and S5-S7)**. Genome wide profiling of H3K4me3 revealed no significant global alterations in promoter associated enrichment across conditions, indicating that transcription start site remained largely preserved under conditions of L1 induced genotoxic stress **(Fig. S5A)**. These data suggest that ORF2p endonuclease activity does not broadly perturb promoter associated chromatin, prompting us to focus on distal regulatory regions. In contrast, H3K27ac profiling revealed extensive redistribution of enhancer associated acetylation in both wild type and p53^-/-^ cells expressing L1 EN⁺ relative to L1 EN⁻ control **(Fig. S5B)**. Pie chart demonstrated increased H3K27ac deposition at distal promoter and intergenic regions in p53^⁻/⁻^ L1 EN⁺ cells relative to their respective input controls **(Fig. S5C and S5D),** indicating that L1 associated DNA damage preferentially coincides with regulatory chromatin. Genome wide classification of H3K27ac marked regions revealed widespread remodelling upon L1 EN activity. In WT cells, we identified gained (n = 5,989), newly gained (n = 6,247), maintained (n = 8,911), and lost (n = 9,186) H3K27ac regions (**Fig. 6C**), whereas in p53^⁻/⁻^ cells we observed gained (n = 9,461), newly gained (n = 8,306), maintained (n = 3,774), and lost (n = 3,418) regions (**Fig. 6D**). Box-and-whisker plots demonstrated increased H3K27ac signal intensity at gained regions, decreased signal at lost regions, and relative stability at maintained regions, highlighting the capacity of ORF2p endonuclease activity to reshape enhancer associated chromatin states. Notably, the magnitude and extent of enhancer remodelling were higher in p53^-/-^ cells, probably due to altered chromatin responses to DNA damage under compromised genome surveillance.

**Figure 6.**
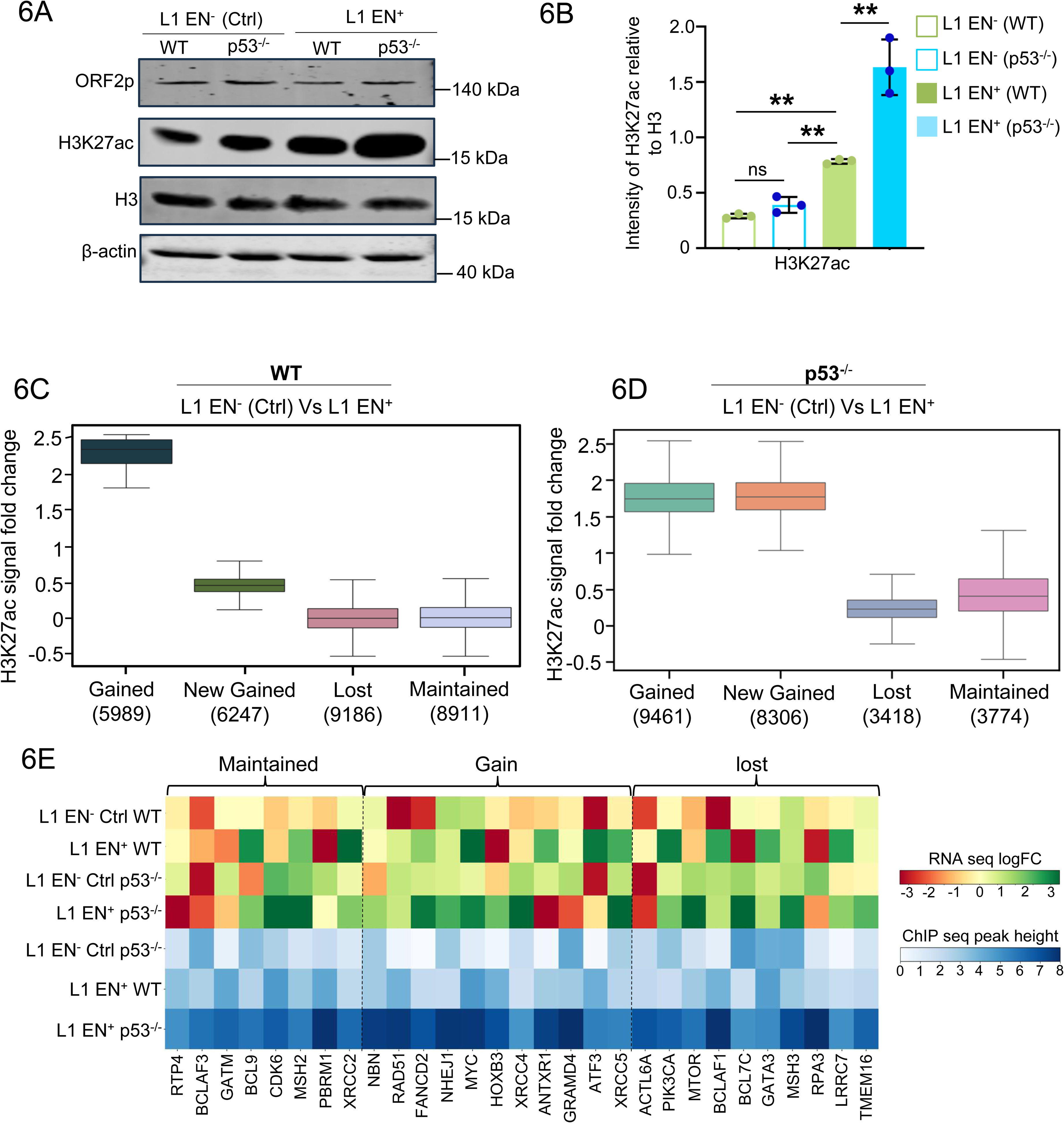
H3K27ac ChIP-seq analyses across WT and p53^-/-^ cells expressing L1 EN^+^ and L1 EN^-^. **(A)** Western blot analysis of H3K27ac histone modification in A375 WT and p53^⁻/⁻^ cells upon L1 EN⁺ or L1 EN⁻ expression. Total H3 was used as a loading control for histone blots, while FLAG immunoblotting confirmed the expression of FLAG tagged ORF2p. β-actin was used as a loading control for whole cell lysates. **(B)** Quantitative analysis of H3K27ac band intensity in A375 WT and p53^⁻/⁻^ cells transfected with L1 EN⁺ (filled) or L1 EN⁻ (unfilled) by using Fiji software. The individual data points in the bar plots represent each biological replicate. The data represents mean ± SD across three biological replicates. Statistical significance was evaluated using the Mann Whitney U test, with significance levels represented as follows: *p<0.05, **p<0.01 and ***p<0.001. **(C)** Box and whisker plot showing the distribution of fold changes in H3K27ac ChIP-seq signal across dynamic regulatory regions upon L1-ORF2p expression. Regions were categorized as new gained (n = 6247), gained (n = 5989), maintained (n = 8911), and lost (n = 9186), based on differential enrichment in L1 EN⁻ versus L1 EN⁺ WT cells. Statistical comparisons using the two-sided Mann Whitney Wilcoxon test revealed significant differences between gained vs. maintained (P = 0.001) and gained vs. lost (P = 0.001) regions, with no significant difference between maintained vs. lost (P = 1.000). Boxes represent interquartile range (IQR), the horizontal line indicates the median, whiskers extend to 1.5× IQR, and individual dots denote outliers. **(D)** Box and whisker plot showing the distribution of fold changes in H3K27ac ChIP-seq signal across dynamic regulatory regions upon L1-ORF2p expression. Regions were categorized as new gained (n = 8,306), gained (n = 9,461), maintained (n = 3,774), and lost (n = 3,418), based on differential enrichment in L1 EN^-^ versus L1 EN^+^ p53^⁻/⁻^ cells. Statistical comparisons using the two-sided Mann Whitney Wilcoxon test revealed significant differences between gained vs. maintained (P = 0.001) and gained vs. lost (P = 0.001) regions, with no significant difference between maintained vs. lost (P = 1.000). Boxes represent interquartile range (IQR), the horizontal line indicates the median, whiskers extend to 1.5× IQR, and individual dots denote outliers. **(E)** Integrated correlation heatmap plot of RNAseq top panel showing relative expression changes in maintained, gain, and lost, and the color codes ranging from red to green, red = down regulation and green = upregulation and the intermediate shade (yellow) = no change. H3K27ac bottom panel, color codes ranges blue shades with high to low normalized peak height. Each column represents genes and rows represent different genotypes.

Together, these data demonstrate that L1-ORF2p endonuclease activity is associated with remodelling of enhancer associated chromatin, particularly at promoter distal and intergenic regions, while leaving promoter regions intact. These effects are amplified in the absence of p53, suggesting that p53 status may influence the chromatin response to EN initiated DNA lesions. To assess whether these chromatin changes were linked to transcriptional outputs, we performed integrative RNA-seq and H3K27ac ChIP-seq analyses in A375 cells expressing L1 EN⁺ or L1 EN⁻. These analyses revealed gained H3K27ac signal at genes involved in DNA repair including NBN, multiple NHEJ pathway components, FANCD2, MYC, and XRCC5, whereas enhancer signal was reduced at loci such as MTOR, BCL7C, and MSH3, and remained stable at XRCC2, BCL9, and GATM **(Fig. 6E)**.

### Endonuclease induced γH2AX accumulation depends on ATM signalling and overlaps with marked H3K27ac chromatin

To determine whether downstream changes observed upon LINE-1 endonuclease (EN) expression are mediated through canonical DNA damage signalling, we inhibited ATM activity, a central kinase in the DNA damage response (DDR), and examined its effect on EN induced yH2AX levels, a marker of DNA damage. A schematic of the experimental workflow illustrated **(Fig. 7A)**. Immunoblot analysis showed that EN expression led to increased ATM phosphorylation (pATM) along with elevated γH2AX levels, indicating activation of DNA damage response signalling. H3K27ac levels were also elevated in EN expressing cells, suggesting that EN induced DNA damage is associated with increased chromatin acetylation. Pharmacological inhibition of ATM attenuated EN induced γH2AX accumulation. This was accompanied by a decrease in H3K27ac levels **(Fig. 7B and C)**, supporting a model in which EN associated chromatin acetylation occurs downstream of ATM dependent DNA damage signalling (75). In contrast, treatment with cisplatin, known to cause genome wide DNA damage, robustly induced pATM and γH2AX but did not lead to a comparable increase in H3K27ac levels. This suggests that EN induced DNA damage elicits a distinct chromatin response that differs from generalized genotoxic stress. Since, we observed increased transcript levels of MYC and NF-κB in RNAseq data **(Fig. 2B and I)**, we performed time-course RT-PCR analysis in EN expressing and cisplatin treated cells. A schematic of the experimental workflow is illustrated in **(Fig. 7D)**. EN expression resulted in a progressive increase in transcript levels of MYC and NF-κB at 24 and 48 hours **(Fig. 7E and F)**. In contrast, cisplatin treatment elicited only modest changes. This observation also suggest that EN induced DNA damage may contribute to oncogenic signalling programs. Next to investigate whether P300 mediates EN-associated chromatin acetylation, we treated cells with a P300 catalytic inhibitor under conditions of EN expression or cisplatin treatment **(Fig. 7G and H)**. Immunoblot analysis revealed a reduction in H3K27ac levels following P300 inhibition in EN-expressing cells, whereas total P300 protein levels remained unchanged. In contrast, cisplatin-treated cells exhibited minimal changes in H3K27ac upon P300 inhibition, further supporting a specific link between EN induced DNA damage and P300 dependent chromatin remodelling.

**Figure 7:**
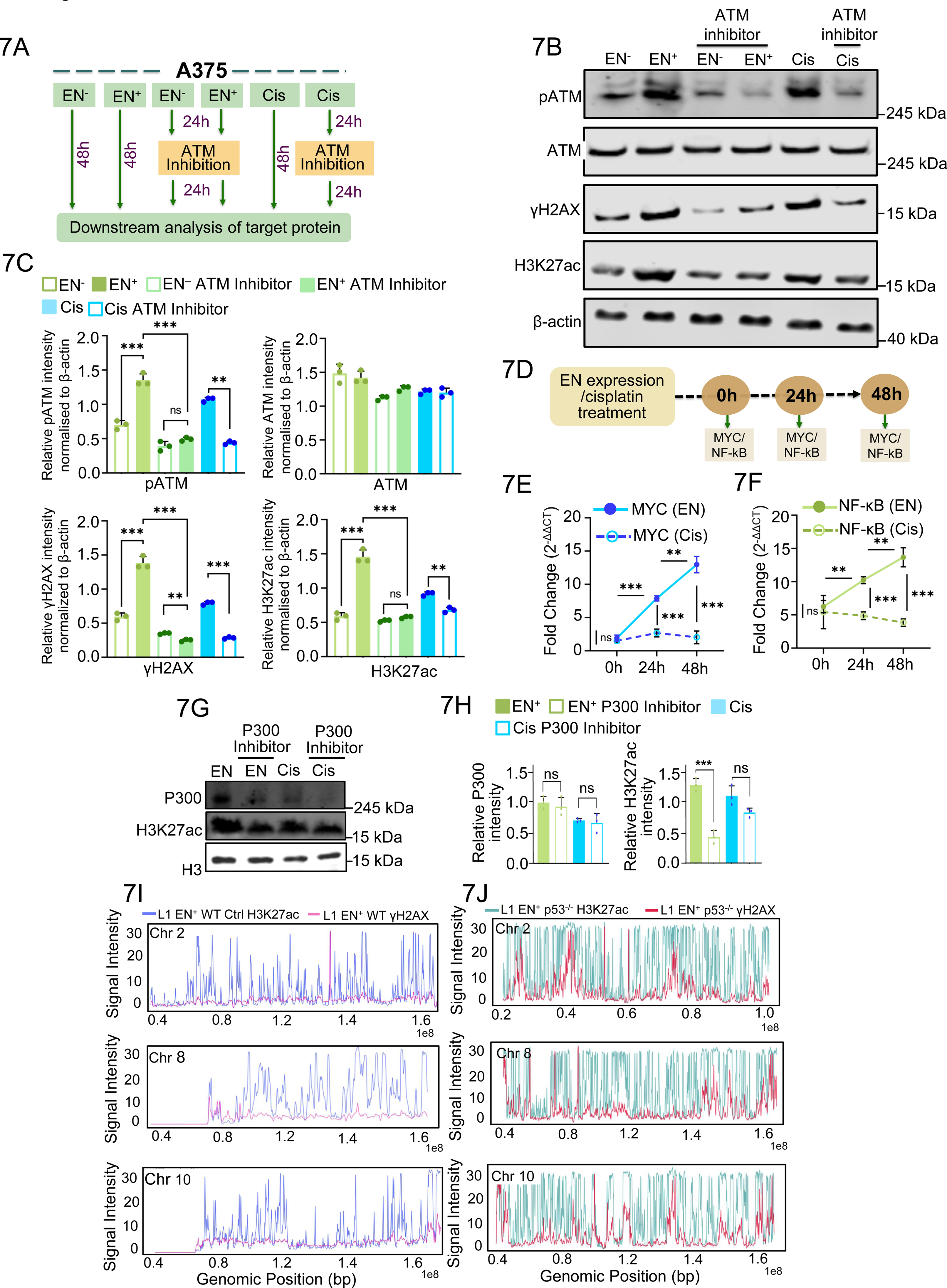
L1 ORF2p endonuclease activity induced DNA damage Signalling and associated changes in H3K27ac levels. **(A)** Schematic overview of the experimental workflow for ATM inhibition studies in A375 cells. DNA damage was induced either by EN expression or cisplatin treatment, followed by ATM inhibition to assess and compare ATM signaling activation under these distinct DNA damage conditions. **(B)** Western blot analysis of phosphorylated ATM (pATM), total ATM, γH2AX, and H3K27ac levels in WT A375 cells expressing EN⁻ or EN⁺. Cells were treated with the ATM inhibitor KU-55933 (10 µM) for 24 hours following EN expression. In parallel, A375 cells were treated with cisplatin (Cis) for 24 hours after ATM inhibition. β-actin was used as a loading control. **(C)** Quantification of western blot band intensities for pATM, total ATM, γH2AX, and H3K27ac in WT A375 cells under EN expression or cisplatin treatment with ATM inhibition. Band intensities were measured by densitometry and normalized to β-actin. Data represent mean ± SD from independent biological replicates, with statistical significance determined using an unpaired Student’s t-test (*p < 0.05, **p < 0.01, ***p < 0.001). **(D)** Schematic illustration of the experimental workflow for assessing MYC and NF-κB expression in EN expressing or cisplatin-treated A375 cells. Samples were collected at 0, 24, and 48 hours for subsequent RT-qPCR analysis. **(E-F)** A375 cells were either transfected with EN⁺ or treated with cisplatin (5 µM) for inducing DNA damage and total RNA was harvested at 0, 24, and 48 hours. RT-qPCR was performed to measure mRNA expression levels of **(E)** MYC and **(F)** NF-κB and values were normalization to β-actin. Data are shown as mean ± SD from three independent biological replicates. Statistical significance was assessed using two-way ANOVA across time points or relative to control (*p < 0.05, **p < 0.01, ***p < 0.001). **(G and H)** To assess activation of P300, DNA damage was caused in A375 either by expressing EN+ for 24 or by subjecting these cells with Cisplatin drug for 24 h. After wards, these cells were treated with P300 inhibitor (3 µM, A-485) for 24 h. Following treatment, cell lysates were subjected to western blot analysis using anti P300 and H3K27ac antibodies. Total histone H3 was used as a loading control. Band intensities were quantified by densitometry and normalized to H3. Data represent three independent biological replicates (n=3) and are presented as mean ± SD. Statistical significance was determined using an unpaired Student’s t test (*p < 0.05, **p < 0.01, ***p < 0.001). **(I)** Genome wide distribution of ChIP-seq signal intensities for H3K27ac (blue) and γH2AX (red) along chromosomes 2, 8 and 10 in L1 EN^+^ p53^⁻/⁻^ A375 cells. Signal values exceeding 30 bin were compressed to enhance visualization of overall peak distribution. The x-axis denotes genomic position (bp), and the y-axis represents normalized signal intensity. **(J)** Genome wide distribution of ChIP-seq signal intensities for H3K27ac (blue) and γH2AX (red) along chromosomes 2, 8 and 10 in L1 EN⁺ WT A375 cells. Signal values exceeding 30 bin were compressed to improve visualization. The x-axis denotes genomic position (bp), and the y-axis represents normalized signal intensity.

**Figure 8:**
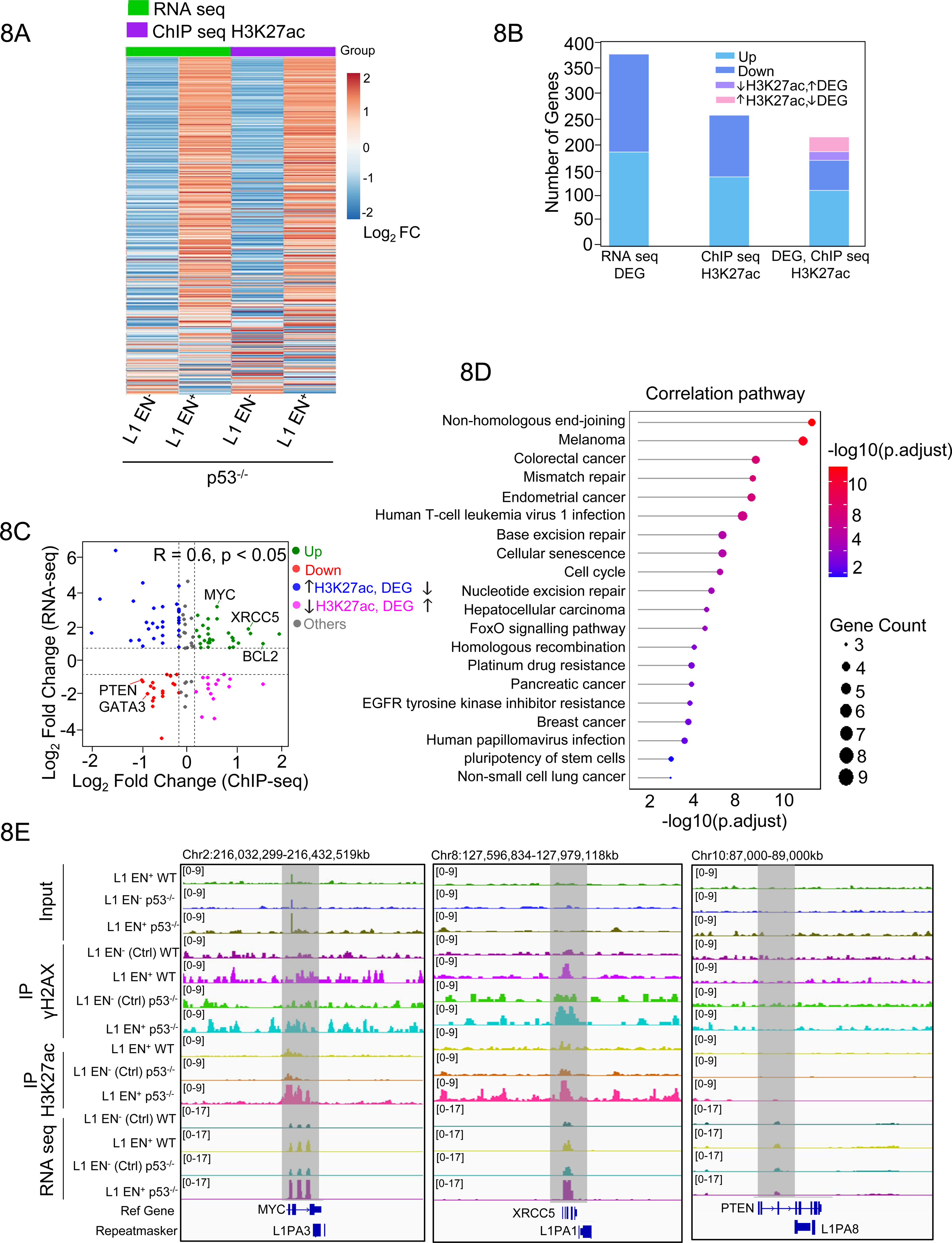
L1 ORF2p endonuclease mediated H3K27ac marked enhancer activation and activation of oncogenic pathways. **(A)** Global heatmap showing differential gene expression and H3K27ac signal in L1 EN⁻ p53^⁻/⁻^ and L1 EN⁺ p53^⁻/⁻^ cells. Hierarchical clustering was performed using RNA-seq (green) and H3K27ac ChIP-seq (purple) data. Each row represents a gene, and columns correspond to biological replicates from L1 EN⁻ p53^⁻/⁻^ and L1 EN⁺ p53^⁻/⁻^ cells. The color scale denotes standardized signal intensities, with red indicating higher mRNA expression or H3K27ac levels, blue indicating lower levels, and white representing intermediate values. **(B)** Stacked bar plot summarizing differential expression and H3K27ac signal changes. RNA-seq analysis identified 193 upregulated and 198 downregulated genes. ChIP-seq analysis revealed 143 upregulated and 124 downregulated H3K27ac marked regions. Among these, 115 genes were common between RNA-seq and ChIP seq datasets and were classified into four categories: upregulated in both datasets (RNA↑ and ChIP↑; n = 22), downregulated in both (RNA↓ and ChIP↓; n = 14), discordant regulation (RNA↑ and ChIP↓; n = 19 and RNA↓ and ChIP↑; n = 12), and other patterns (n = 48) **(C)** Scatter plot illustrating concordant and discordant changes in gene expression and H3K27ac occupancy. Each point represents a gene plotted by log₂ fold change in RNA-seq (y axis) versus H3K27ac ChIP-seq (X axis). Genes showing concordant upregulation (green) or downregulation (red) suggest enhancer linked transcriptional control. Discordant genes include those upregulated in RNA-seq but downregulated in H3K27ac (blue), or vice versa (magenta). Grey dots represent genes with non-significant or inconsistent changes. Notable examples include MYC, BCL2, and XRCC5 (concordantly upregulated), and PTEN and GATA3 (concordantly downregulated). **(D)** Lollipop plot showing enriched KEGG pathways among genes associated with regions gaining H3K27ac signal. The X axis represents the gene ratio (number of input genes in the pathway divided by the total number of pathway associated genes), and the y axis lists the pathway names. Dot size reflects the number of genes, and color indicates adjusted p values, with darker red representing higher statistical significance. **(E)** IGV snapshots of XRCC5, MYC and PTEN loci across L1 EN⁻ and L1 EN⁺ conditions in WT and p53^⁻/⁻^ A375 cells. Tracks display γH2AX, H3K27ac ChIP-seq and RNA-seq signals. Shaded areas of XRCC5, MYC mark loci where elevated H3K27ac signal aligns with increased transcript levels, whereas the shaded areas of PTEN exhibits decreased γH2AX, H3K27ac signal conciding with decreased transcript levels.

To determine whether ORF2p endonuclease induced DNA damage occurs preferentially within specific chromatin contexts, we performed γH2AX ChIP-seq in A375 WT and p53^⁻/⁻^ cells expressing L1 EN⁺ or EN⁻ **(Fig. S6A).** Genome-wide analyses revealed a redistribution of γH2AX signal toward distal promoter and intergenic regions in p53^⁻/⁻^ EN⁺ cells, compared to controls (**Fig. S6B-E**), suggesting preferential accumulation of DNA damage within transcriptionally active regulatory regions. Integration of γH2AX and H3K27ac ChIP-seq datasets demonstrated spatial concordance between DNA damage sites and active chromatin regions. Genome-wide signal distribution revealed substantial overlap between γH2AX-enriched regions and H3K27ac-marked chromatin domains. Analysis of representative chromosomes (2, 8, 10, 15, 17, and 18) further confirmed increased co-enrichment of γH2AX and H3K27ac signals in EN⁺ conditions in both WT and p53^-/-^ cells compared to EN⁻ controls (**Figs. 7I and J S6F and G)**.

For visualization purposes, signal intensities exceeding 30 bins were capped to facilitate comparison of global distribution patterns. Collectively, these findings demonstrate that ORF2p endonuclease activity induces focal γH2AX accumulation that spatially coincides with H3K27ac enriched chromatin. Loss of p53 further amplifies this effect, suggesting a model in which LINE-1 activity links DNA damage signalling to epigenomic remodelling within regulatory chromatin landscapes.

### L1-ORF2p endonuclease induces enhancer and oncogenic transcriptional activation

To determine whether ORF2p endonuclease induced DNA damage is accompanied by alterations in enhancer activity and gene expression, we integrated H3K27ac ChIP-seq with RNA-seq datasets from A375 p53^⁻/⁻^ expressing L1 EN⁺ or L1 EN⁻. Global heatmap analysis revealed distinct patterns of gene activation and repression between EN⁺ and EN⁻ conditions across both H3K27ac ChIP-seq and RNA-seq datasets, indicating a strong association between ORF2p endonuclease activity, remodelling of the enhancer landscape, and altered transcriptional output **(Fig. 8A)**. Transcriptomic analysis of A375 p53^⁻/⁻^ cells expressing L1 EN⁺ identified, 391 differentially expressed genes (DEGs 193 upregulated, 198 downregulated), while H3K27ac ChIP-seq revealed 267 regions with altered acetylation (143 gained and 124 lost) relative to L1 EN⁻. Integration of RNA-seq and H3K27ac datasets identified 115 genes exhibiting concordant transcriptional and enhancer changes, indicating coordinated epigenetic transcriptional regulation downstream of ORF2p endonuclease activity. Of these, 22 genes showed concurrent transcript upregulation and H3K27ac gain, whereas 14 displayed coordinated downregulation. In contrast, 12 genes were transcriptionally repressed despite enhancer gain, and 19 genes lost H3K27ac without detectable changes in mRNA levels, indicating partial uncoupling between enhancer acetylation and mRNA levels **(Fig. 8B)**.

To assess the influence of p53, we performed similar analysis in L1 EN⁺ expressing WT A375 cells, identifying 757 DEGs (560 upregulated, 197 downregulated) and 386 differentially acetylated H3K27ac regions (179 gained, 207 lost). Integration of these datasets revealed 281 genes with coordinated epigenetic and transcriptional changes, demonstrating that ORF2p endonuclease activity elicits a broad enhancer associated transcriptional responses. Among these, 103 genes showed transcript upregulation and H3K27ac gain, whereas 78 exhibited coordinated loss. Additional subsets displayed discordant regulation, including transcript downregulation with enhancer gain (48 genes) and enhancer loss without measurable transcript change (34 genes), highlighting context dependent regulatory effects **(Fig. S7A)**.

To quantitatively assess enhancer transcript relation, we plotted gene wise log₂ fold changes in RNA-seq against corresponding changes in H3K27ac signal in both p53^⁻/⁻^ and WT A375 cells **(Figs. 8C and S7B)**. A subset of genes, including oncogenes such as MYC, BCL2, and XRCC5, exhibited concordant upregulation at both transcriptional and enhancer levels (Up-Both), whereas tumor suppressors such as PTEN and GATA3 showed coordinated downregulation (DownBoth). These patterns indicate selective, enhancer linked transcriptional regulation rather than global activation or repression, consistent with targeted chromatin remodelling.

Motif enrichment analysis of regions with gained or lost H3K27ac revealed significant enrichment of transcription factor binding motifs, including multiple KRAB-ZNF motifs within newly gained enhancers **(Fig. S7C-S7E)**. Given the established role of KRAB-ZNF proteins in transposable element repression, this enrichment suggests that ORF2p activity, particularly in the context of p53 loss, may derepress transposon-associated regulatory sequences, thereby exposing latent enhancer elements that subsequently acquire H3K27ac.

Gene set enrichment analysis (GSEA) of genes associated with gained H3K27ac regions in p53^⁻/⁻^ EN⁺ cells identified enrichment of pathways related to cancer signalling and DNA damage response **(Fig. 8D)**, consistent with notion that genotoxic stress can reprogram enhancer landscapes to support adaptive transcriptional programs. Visualization of representative loci using IGV, including XRCC5, MYC, BCL2, HOXB13, and RepeatMasker annotated regions, demonstrated concordant enrichment of γH2AX, H3K27ac, and RNAseq signal in EN⁺ cells **(Figs. 8E and S7F)**. Collectively, these findings demonstrate that ORF2p endonuclease activity drives focal enhancer remodelling and selective transcriptional rewiring. Loss of p53 further amplifies the extent and persistence of these enhancer associated regulatory changes, highlighting a mechanism by which LINE-1 activity links DNA damage signalling to epigenomic reprogramming and oncogenic transcriptional activation.

## Discussion

Our findings support a model in which the endonuclease activity of LINE-1 ORF2p acts as an initiating trigger of DNA damage signalling, engaging ATM-dependent pathways and subsequently promoting chromatin remodelling. Notably, ORF2p-mediated DNA cleavage appears to preferentially occur at fragile genomic regions. Consistent with this, EN⁺ induced DNA damage led to enhanced deposition of γH2AX at these sites, as confirmed by ChIP-qPCR, indicating localized DNA damage **(Fig. 1)**.

In contrast, treatment with cisplatin, which induces widespread DNA damage, resulted in robust global γH2AX accumulation but did not show comparable enrichment at these regions **(Fig. S1F)**. This suggests that the chromatin changes observed under EN⁺ conditions are not merely a consequence of global DNA damage signalling, but are associated with regionally biased DNA damage.

Together, these results indicate that ORF2p induced DNA damage may contribute shaping the local epigenomic landscape. However, further investigation is required to distinguish between direct mechanistic coupling and indirect downstream effects of DNA damage signalling, particularly given the broad role of ATM.

The mutagenic potential of LINE-1 insertions is well established (6,76), our results indicate that the oncogenic impact of LINE-1 activity is also driven by ORF2p endonuclease mediated DNA damage signalling activation **Figs. 1-3**. Catalytically EN^-^inactive ORF2p produced minimal detectable activation of DNA damage signalling whereas endonuclease competent ORF2p expression resulted in extensive double strand breaks, ATM dependent γH2AX deposition. These effects were consistently more pronounced in p53^-/-^ cells, supporting a role for p53 in constraining the downstream consequences of ORF2p initiated damage. By employing inhibitor studies, we demonstrate that ATM activation represents an early event in the ORF2p induced DNA damage response, that follows increase in P300 and H3K27ac **(Figs**. **4 and 5)**, Chromatin fractionation experiments show increased association of P300 with the chromatin fraction, accompanied by elevated H3K27ac levels under EN^+^ conditions **(Fig. 5E)**, consistent with activation of enhancer associated chromatin states. Temporal analyses further indicate that γH2AX accumulation precedes recruitment of DNA repair factors such as 53BP1 and XRCC5, followed by H3K27ac enrichment **(Fig. 5B)**, supporting the interpretation that histone acetylation occurs downstream of the DNA damage response. However, our results do not establish direct recruitment of P300 to γH2AX marked loci or a direct mechanistic coupling between DNA damage and P300 activity.

ORF2p induced DNA lesions were associated with a shift towards error prone, NHEJ pathway, as reflected by increased expression and co-localisation of NHEJ components. We think that NHEJ factors are probably preferably recruited to ORF2p induced lesions for damage resolution however, it doesn’t rule out the possibility of the involvement of other repair pathways. Despite widespread enhancer disruption, promoter associated H3K4me3 marks remained stable, revealing a selective vulnerability of enhancer architecture to ORF2p induced genotoxic stress. The emergence of new H3K27ac enriched enhancers coincided with transcriptional activation of oncogenic pathway **(Figs. 6-8)**, consistent with models in which DNA damage signalling intersects with regulatory element activation under conditions of replication stress or inflammation (77). Notably, expression of endonuclease domain constructs recapitulated key aspects of the DNA damage, and transcriptional phenotypes observed with full length ORF2p, establishing that endonuclease activity is able to drive these effects. Upon perturbation of XRCC5 in cancer cells with derepressed LINE-1, we observed a marked increase in DNA damage that led to enhanced cell lethality **(Fig. 7)**. Previous studies have proposed that endogenous LINE-1 activity marks tumors that are sensitive to genotoxic therapies, including inhibition of DNA damage response pathways (78). However, this study directly interrogates ORF2p enzymatic activity or chromatin consequences by directly isolating the contribution of ORF2p endonuclease activity. Within this framework, ATM mediated DNA damage signalling emerges as a key upstream event linking ORF2p induced DNA breaks to P300 recruitment and increased H3K27ac. However, more experiments are required to establish if this is a direct consequential link. We think that in enhancer rich and transcriptionally active regions, ORF2p induced DNA breaks may encounter chromatin environments that favour rapid but error prone repair. Over time, this possible connection of DNA damage and enhancer remodelling may promote transcriptional plasticity, genomic instability, and adaptive gene expression programs that may contribute to tumorigenesis.

In summary, this study identifies LINE-1 ORF2p endonuclease activity as a potent driver of DNA damage signalling that is followed by chromatin associated changes **(Fig. 9)**. Our findings establish a mechanistic link between transposon reactivation, ATM-mediated DNA damage responses, and downstream chromatin modulation in cancer.

### Experimental procedures Cell culturing

p53 wild-type (WT) and p53 knockout (p53^-/-^) cells of A375 and U2OS were obtained from Prof. John Abrams, UT Southwestern Medical Center, Dallas, Texas. A375 cells were cultured in Dulbecco’s Modified Eagle’s Medium (Gibco, #12100046). All cell lines were grown at 37°C, 5% CO_2_ and passaged using Trypsin (Gibco, #15400054) in the presence of an anti-antimycotic antibiotic (Gibco, #15240062).

### Plasmid Constructs

Plasmids pMT646 and pMT1093 (were a kind gift from Prof. Kathleen Burns, Harvard Medical Centre, USA). The pLIX401-APE1 WT construct (Addgene plasmid #227190) was obtained from Addgene.

### Generation of quasi stable cell lines of pMT646 and pMT1093

The pMT646 and pMT1093 constructs are based on the pCEP4 Puro backbone and were introduced into A375 WT and p53^-/-^ cells by transfection. Following transfection, cells were subjected to selection with Puromycin at 2 µg/mL to enrich for populations maintaining episomal plasmids. Puromycin resistant populations were maintained under continuous selection throughout all experiments performed. Expression of L1-ORF1p was validated using an

ORF1p specific antibody, whereas L1-ORF2p expression was assessed in Puromycin selected populations by immunofluorescence and western blotting using anti FLAG antibodies (Invitrogen, #MA191878), since all constructs encode FLAG tagged ORF2p. Both full length L1 and endonuclease specific constructs were assessed where indicated. All experiments shown in **(Fig. 1-4)** were performed in parallel with appropriate controls, including empty vector transfected cells processed under identical conditions. For assays involving transient expression, analyses were conducted within 24 to 48 hours following transfection. All quantitative experiments were performed using a minimum of three independent biological replicates per condition. For imaging, multiple fields of view were analysed per replicate and pooled for quantification. Western blot, immunofluorescence, and chromatin immunoprecipitation experiments were performed using samples from three independent biological replicates to ensure reproducibility.

### Cloning and expression of EN-ORFeus

The EN ORFeus construct was generated by PCR amplification of the endonuclease (EN) domain of L1-ORF2p from pMT646 plasmid. The PCR product was digested with EcoRI and BamHI (New England Biolabs, EcoRI #R3101S, BamHI #R3142S) and cloned in frame into the pEGFP-N3 vector to generate a C-terminal GFP fusion protein under the control of the CMV promoter. To generate the 5′UTR EN-ORFeus construct, the CMV promoter in pEGFP-N3 was replaced with the LINE-1 5′ untranslated region (5′UTR) promoter. The ORF2p endonuclease domain was amplified from pMT646 and cloned into the modified pEGFP-N3 vector using EcoRI and BamHI restriction sites, yielding a GFP tagged EN-ORFeus construct. All constructs were verified by restriction digestion. Primer sequences used for amplification are listed in **Supporting Table 1**. Experiments were performed under transient transfection conditions, and analyses were conducted 24-48 h post transfection.

### Real-Time PCR experiments (qPCR)

Total RNA was isolated using TRIzol (Invitrogen), then treated with Turbo DNase (Thermo Fisher, #AM1907). mRNA was reverse transcribed into cDNA using cDNA synthesis kit Verso (Thermo Fisher, #AB1453A). To carry out qRT PCR experiments 20ng cDNA was used per reaction. Gene specific primers against XRCC5, RAD51, XRCC2, LIG4, MRE11 were used for PCR reaction (primer sequences listed in **Supporting Table 1**). Applied biosciences QuantStudio 5 and QuantStudio 6 were used for real time PCR and normalization of gene expression was done against β-actin (for standard gene expression analysis Fold Change = 2^- ΔΔCT^, ΔΔCT= Δ Target Sample-Δ Reference Sample).

### Chromatin immunoprecipitation

1 × 10^7^ A375 WT and p53^-/-^ cells stably transfected with pMT646 (L1 EN^+^), pMT1093 (L1 EN^-^) were grown in a 150 cm^2^ dish and were fixed with 0.75% formaldehyde in dish at room temperature for 10 minutes while gentle shaking and subsequently quenched by 125mM glycine solution was added to stop the over fixation. The cells were scraped from the dish and were resuspended in ice cold ChIP lysis buffer (50 mM HEPES-KOH pH 7.5, 140 mM NaCl, 1 mM EDTA pH 8, 1% Triton X-100, 0.1% Sodium deoxycholate, Protease inhibitor (Sigma, #539131). The cell lysate (i.e., chromatin) was sonicated using an M220 Focused-ultrasonicator (Covaris) to fragment chromatin lengths of 400 bp (peak power 75.0, Duty factor = 10, Cycles/Burst = 200, for 240 seconds). For each immunoprecipitation 25μg chromatin was diluted ten times with RIPA buffer and incubated with the following antibodies: γH2AX (Active motif, #39117), H3K27ac (Cell Signalling Technologies, #8173), H3K4me3 (Cell Signalling Technologies, #9751). Agarose A/G beads (EMD Millipore, #IP0515ML), were charged and incubated with the antibodies overnight, following low salt, high salt, and LiCl washes the next day and eluted. Eluted chromatin was treated with RNAse A (Thermo Fisher, #12091021) and proteinase K (NEB, #P8107S), crosslinks were reversed, and samples were purified using a PCR purification kit (Qiagen). ChIP enrichment was quantified with active histone mark using β-Actin by qRT-PCR with QuantStudio 5 and QuantStudio 6.

### ChIP-sequencing

After Immunoprecipitation indexed NGS libraries were prepared by end repairing, A tailing, adaptor ligation, and amplification procedures according to manufacturer’s instructions in NEBNext® Ultra™ II DNA Prep Kit (New England Biolabs, #E7645,). Upon library preparation, libraries were analysed using Agilent Technologies 4200 Tapestation followed by sequencing in Illumina NovaSeqTM 6000 with 150 bp paired end reads.

### Chromatin immunoprecipitation-qPCR

A375 and U2OS WT and p53^-/-^ cells expressing either pMT646 (L1 EN⁺) or pMT1093 (L1 EN⁻) were generated as quasi stable pooled populations by selection and maintenance in Puromycin. Cells were harvested and chromatin was cross linked prior to immunoprecipitation with an anti γH2AX antibody (Active Motif, #39117) following the ChIP protocol described above. Immunoprecipitated chromatin was eluted in nuclease free water, diluted tenfold, and analysed by qPCR using primers targeting fragile site hotspots at FRA3B (FCR and FDR regions). ACTB was used as a negative control. In parallel, the same ChIP-qPCR experiment was performed in cells transfected with the LINE-1 EN catalytic domain alone, with comparison to cells treated with cisplatin (5 μM) as a positive control for DNA damage. Primer sequences are provided in **Supporting Table 1**.

### CRISPR Cas9 mediated XRCC5 depletion and L1 endonuclease expression

A single guide RNA (sgRNA) targeting XRCC5 (forward: 5′-CACCGTCATAAGGCATATTCGACTA-3′; reverse: 5′-AAACTAGTCGAATATGCCTTATGAC-3′), adopted from the Feng Zhang laboratory, was cloned into the pTLCV2 CRISPR Cas9 vector (A kind gift from Dr. Vinay Bulusu, IISER Berhampur) using the BsmBI restriction enzyme (NEB, #R0580). Correct insertion of the sgRNA was confirmed by Sanger sequencing.

WT A375 cells were co-transfected with CRISPR-Cas9 plasmid carrying sgRNA targeting XRCC5 and L1-EN⁺ or L1-EN^-^. 48 hours after transfection, cells were washed with ice cold phosphate buffered saline (PBS) and lysed in RIPA buffer supplemented with protease and phosphatase inhibitors. Cell lysates were centrifuged at 12,000 × g for 15 min at 4°C, and the supernatant was obtained was subjected to bicinchoninic acid (BCA) assay for protein estimation. Equal amounts of protein were resolved by SDS-PAGE and transferred to nitrocellulose membrane. Membrane was blocked with 3% (w/v) non fat dry milk in PBS containing 0.1% Tween 20 (PBST) and incubated overnight at 4°C with primary antibodies against XRCC5 (Ku80), γH2AX, and β-actin. After incubation with appropriate fluorescently conjugated secondary antibodies, signals were detected using a LICOR (ODYSSEY) fluorescence imaging system. β-actin was used as a loading control for all immunoblot analyses.

### MTT Assay

10,000 A375 WT cells were seeded into 96-well plates a day prior to transfection to achieve approximately 60-70% confluence at the time of transfection. Cells were co-transfected with a CRISPR-Cas9 sgRNA targeting XRCC5 and L1 EN⁺. Cells transfected with either with pTLCV2 CRISPR-Cas9 only or L1 EN⁺ served as negative controls. Each experimental condition was plated in at least three technical replicates per experiment, and all experiments were independently repeated at least three times (biological replicates). 48 hours post transfection; cell viability was assessed using an MTT assay. Briefly, MTT reagent was freshly prepared in sterile PBS and added to each well to a final concentration of 0.5 mg/ml. Plates were incubated at 37°C for 3-4 hours in the dark to allow the formation of insoluble formazan crystals. The culture medium was then carefully removed without disturbing the crystals, and formazan was solubilized by the addition of dimethyl sulfoxide. Plates were gently agitated on a plate shaker for 5-10 min until complete dissolution was achieved. Absorbance was measured at 570 nm using a microplate reader (SpectraMax® iD5 Multi-Mode Microplate Reader), with background correction performed using a reference wavelength between 630 and 690 nm where available. Blank wells containing medium and MTT reagent, but no cells were included for background subtraction. Cell viability was calculated as a percentage relative to the appropriate control condition following blank correction.

### Immunofluorescence

Quasi stable cell lines of A375 WT and p53^-/-^ cells expressing pMT646 or pMT1093 were cultured on poly L lysine coated coverslips (Sigma, #P4707). Cells were washed with 1X PBS and fixed in 4% paraformaldehyde for 15 minutes at room temperature. Following fixation, cells were permeabilized with 0.1% TritonX-100 in 1X PBS for 10 minutes and blocked with 1% fetal bovine serum in 1X PBST (0.1 percent Tween-20 in PBS) for 30 minutes at room temperature. Primary antibodies used were FLAG (Invitrogen, #MA191878), γH2AX (Active Motif, #39117), XRCC5 (Cell Signalling Technology, #2180), and RAD51 (Cell Signalling Technology, #65653). Secondary antibodies included anti-mouse 568 (Thermo Fisher, #A11031), anti-mouse 488 (Thermo Fisher, #A11001), anti-rabbit 488 (Thermo Fisher, #A10680), and anti-rabbit 568 (Thermo Fisher, #A 11011), all used at a dilution of 1:1000.

For experiments using CMV driven EN-GFP and L1 5′ UTR driven EN-GFP constructs, A375 WT p53^⁻/⁻^ cells were transiently transfected under identical conditions. Cells were seeded onto poly-L-lysine coated glass coverslips to ensure uniform attachment and optimal imaging quality. Cells were fixed and analysed 24 h post transfection. Following fixation, cells were processed for immunofluorescence staining using the same antibodies, staining protocols, and wash conditions described above. GFP fluorescence was used to identify EN expressing cells, and γH2AX immunostaining was employed as a marker of DNA damage. Image acquisition was performed using identical microscope settings (laser power, exposure time, and detector gain) across all conditions. This standardized imaging approach enabled direct qualitative comparison of DNA damage induced by CMV driven expression versus native L1 5′ UTR driven EN expression.

### Pharmacological Inhibition of ATM, P300 followed by Immunoblot Analysis

A375 cells were transfected with LINE-1 endonuclease or treated with cisplatin (5 µM) under the indicated experimental conditions. To inhibit ATM kinase activity, cells were treated with the selective ATM inhibitor KU-55933 (Medchem, #HY-12016) (10 µM) for 24 hours following transfection and/or cisplatin treatment. For inhibition of P300 catalytic activity, cells were treated with the selective P300 inhibitor A-485 (Medchem, #HY-107455) (3 µM) for 48 hours under EN expressing or cisplatin treated conditions. For whole cell protein analysis, cells were lysed in ice-cold RIPA buffer (50 mM Tris-HCl, pH 7.4, 150 mM NaCl, 1% NP-40, 0.5% sodium deoxycholate, and 0.1% SDS) supplemented with protease and phosphatase inhibitors. Lysates were incubated on ice for 20-30 min with intermittent mixing and clarified by centrifugation at 13000 rpm for 30 min at 4°C. Protein concentrations were determined using a BCA assay, and equal amounts of protein were resolved by SDS-PAGE and transferred onto nitrocellulose membranes. Membranes were probed with primary antibodies against Total ATM (Cell Signaling Technologies, #2873), Phosphorylated ATM (pATM) (Cell Signaling Technologies, #13050), γH2AX (Active motif, #39117), H3K27ac (Cell Signaling Technologies, #8173), and P300 (Cell Signaling Technologies, #54062), with β-actin (Merck Millipore, #A2228) or histone H3 (Cell Signaling Technology, #14269) used as loading controls as appropriate. This was followed by incubation with species-specific DyLight 800-conjugated secondary antibodies (Cell Signaling Technologies, #5257, #5151) (1: 10,000 dilutions in PBS containing 3% BSA). Protein signals were detected using the LI-COR Odyssey imaging system. Densitometry analysis was performed using ImageJ (or LI-COR software), and protein levels were normalized to their respective loading controls.

### Subcellular Fractionation and Western Blot Analysis

A375 cells were harvested under the indicated experimental conditions and subjected to biochemical fractionation as previously described with minor modifications (79,80). Briefly, cells were washed twice with ice-cold PBS and lysed in hypotonic lysis buffer (10 mM HEPES, pH 7.9, 10 mM KCl, 1.5 mM MgCl₂, 0.1% NP-40) supplemented with protease and phosphatase inhibitors to isolate the cytoplasmic fraction. Lysates were incubated on ice for 10 min and centrifuged at 1,000 × g for 5 min at 4°C. The supernatant was collected as the cytoplasmic fraction. The nuclear pellet was washed and resuspended in nuclear extraction buffer (20 mM HEPES, pH 7.9, 400 mM NaCl, 1 mM EDTA, supplemented with protease and phosphatase inhibitors) and incubated on ice for 20-30 min with intermittent mixing to extract nuclear soluble proteins. Following centrifugation at 14,000 × g for 10 min at 4°C, the supernatant was collected as the nuclear soluble fraction. The remaining insoluble pellet, enriched for chromatin associated proteins, was washed and subjected to chromatin solubilisation using SDS containing buffer (0.5%) along with DNase I treatment, followed by sonication to ensure efficient extraction of chromatin bound proteins. This fraction was collected as the chromatin associated fraction. Protein concentrations were determined using a BCA assay, and equal amounts of protein from each fraction were resolved by SDS-PAGE and transferred onto nitrocellulose membranes (Bio-Rad, #1620112). Membranes were probed with primary antibodies against XRCC5 (Ku80) (Cell Signaling Technology, #2180), P300 (Cell Signaling Technologies, #54062), H3K27ac (Cell Signaling Technologies, #8173), γH2AX (Active motif, #39117), total H3 (Cell Signaling Technology, #14269), and β-actin (Merck Millipore, #A2228), followed by species-specific DyLight 800 conjugated secondary antibodies (Cell Signaling Technologies, #5257, #5151) (1: 10,000 dilution in PBS containing 3% BSA). Fluorescent signals were detected using the LI-COR Odyssey imaging system. β-actin was used as a cytoplasmic loading control, while histone H3 served as a chromatin fraction control.

### RNA-sequencing

Total RNA was extracted using TRIzol reagent (Thermo Fisher Scientific) from L1 EN^+^ WT, L1 EN^+^ p53^-/-^, L1 EN^-^ WT, L1 EN^-^ p53^-/-^ in A375 melanoma cell lines. RNA integrity was checked by gel electrophoresis, the library preparation of total RNA performed according to manufacturer’s instructions, NEBNext® Ultra™ II RNA Prep Kit for illumina (Cat. #E7775, NEB). The quality of library was analysed using Agilent Technologies 4200 Tapestation and sequenced in Illumina NovaSeqTM 6000 with 150 bp paired end reads.

### Microscopy

Images were captured using 60X oil objective Leica STELLARIS 8 Falcon and ZEISS Apotome 3 (IISER Berhampur). All images were processed via Fiji software package and the bioinformatics imaging quantification pipeline.

### Microscopy Quantification

Digital images of GFP positive cells were used to quantify the total number of γH2AX foci per nucleus. For each biological replicate, a minimum of 30 non-overlapping fields of view were acquired, corresponding to approximately 100-150 nuclei per condition per replicate. Quantification was performed across three independent biological replicates, yielding a total of approximately 300-450 nuclei analysed per condition. Image analysis was carried out using ImageJ (Fiji) with a standardized and blinded analysis pipeline applied uniformly across all conditions. Briefly, background subtraction and thresholding were performed prior to foci detection. γH2AX foci were identified using a size filter of 0.5-1.5 µm² to capture discrete DNA damage foci; larger γH2AX domains (>1.5 µm²), which likely represent clustered or pan nuclear damage, were excluded from foci number quantification but were retained for qualitative assessment. The same analysis parameters were applied to XRCC5 and RAD51 foci quantification to ensure consistency across markers. For colocalization analysis, only nuclei with clearly resolved foci and minimal overlap were included to reduce artefacts arising from densely damaged regions. Colocalization between γH2AX and XRCC5 or RAD51 was quantified using Pearson correlation coefficients calculated on a per nucleus basis, with coefficients between

0.6 and 1.0 considered indicative of significant colocalization. Colocalization metrics were averaged across all analysed nuclei within each replicate and then across biological replicates. Statistical analyses were performed using the Mann-Whitney U test unless otherwise stated. Significance thresholds were defined as *p < 0.05, **p < 0.01, and ***p < 0.001. All quantifications were performed using identical image acquisition settings and analysis parameters across experimental conditions to enable direct comparison.

### Alkaline and neutral COMET assay

The Empty Vector (control), EN-GFP transfected positive cells were extracted and comet assays were performed. Alkali electrophoresis buffer consisted of 200 mM NaOH, 1 mM EDTA and pH 13. Neutral electrophoresis buffer (TBE buffer) consisted of 89 mM Tris-base, 89 mM Boric acid, 2 mM EDTA and pH 8.3. The slides were then stained with PI (7.5ug/ml) for 10 min and images were visualized under a PI filter (535 nm) with an Apotome microscope (ZEISS Apotome 3). The comets were analysed using ImageJ image analysis software. A total of 5 field comet images from each sample of each biological replicate were evaluated. Statistical significance was evaluated using the Mann-Whitney U test, with significance levels represented as follows: *p < 0.05.

### Western Blot

Protein extraction was performed from A375 WT and p53^-/-^ quasi stable cells expressing L1 EN⁺ or catalytically inactive L1 EN⁻ constructs. Untransfected parental A375 WT and p53^-/-^cells processed in parallel served as baseline controls. Cells were harvested at identical time points and lysed on ice using RIPA buffer supplemented with a protease inhibitor cocktail (Sigma, #539131) to minimize proteolytic degradation. Lysates were centrifuged at 14,000 × g for 15 minutes at 4 °C, and the supernatant was used to estimate protein concentration using bicinchoninic acid (BCA) assay (Thermo Fisher Scientific, #23227). To ensure quantitative accuracy and reproducibility, sample loading conditions were optimized by performing preliminary titration experiments to identify the linear dynamic range for each antibody. Equal amounts of total protein, selected based on these optimization experiments, were resolved by SDS-PAGE and transferred onto nitrocellulose membrane (Bio-Rad, #1620112). Membrane was blocked for 1 hour at room temperature in phosphate buffered saline containing 0.1% Tween-20 (PBST) and 3% non-fat dry milk, followed by incubation with primary antibodies diluted 1:1000 in 1× PBS containing 3% bovine serum albumin (BSA). Primary antibody incubations were performed overnight at 4°C with gentle agitation. The following antibodies were used: γH2AX (Active Motif, #39117), XRCC5/Ku80 (Cell Signaling Technology, #2180), H3K27ac (Cell Signaling Technology, #8173), FLAG tag (Invitrogen, #MA1-91878), β-actin (Merck Millipore, #A2228), and histone H3 (Cell Signaling Technology, #14269).

After washing in PBST, membranes were incubated with species specific anti-mouse or anti-rabbit IgG (H+L) secondary antibodies conjugated to DyLight 800 (4× PEG; diluted 1: 10,000 in 1× PBS containing 3% BSA). Fluorescent signals were detected using LICOR (ODYSSEY) imaging system.

γH2AX signal intensities were quantified and normalized to FLAG tagged ORF2p expression to specifically assess L1-ORF2p associated DNA damage, thereby minimizing confounding effects arising from variable transfection efficiencies. β-actin and histone H3 were used as loading controls for whole cell lysates. Densitometry analyses were performed using identical processing parameters across all conditions. All experiments have been performed 3 times and representative results of one experiment are shown.

We used Fiji software to quantify the Western blot signals. First, we selected the lanes corresponding to each sample using the rectangle tool and then used the "Plot Lanes" function to generate intensity profiles of the bands. The area under each peak, representing the band intensity, was measured. To ensure accurate comparisons, we calculated the percentage contribution of each band to the total signal and normalized the target protein bands to the total protein loaded. Finally, the values were expressed relative to the control sample, which was set to 1.

### Statistics

Statistical analyses were performed using Prism 7.01 (GraphPad Inc.). For comparisons between two groups, either a two-tailed Student’s t test or a nonparametric Mann-Whitney U test was used, as indicated in the figure legends, depending on data distribution. For multiple group comparisons, one-way or two-way analysis of variance (ANOVA) with appropriate post hoc correction was applied. A p value < 0.05 was considered statistically significant.

### RNA-seq data analysis

Adaptor sequences and low quality bases were trimmed from raw sequencing reads using FASTP (81) (0.32.2). Reads shorter than 30 base pairs after trimming were discarded. Quality control of raw reads was assessed using FASTQC (https://github.com/s-andrews/FastQC) (v0.11.9), and summary reports were inspected to confirm read quality, base composition, and adapter removal efficiency prior to downstream analyses.

### Gene expression analysis

The remaining clean set of reads were then aligned to the reference genome build hg38 (GRCh38) with STAR (82) (2.7.10a) using the default parameters. Analysis of the repeat genome was performed separately (see below for details). Gene expression levels were estimated by quantifying primary aligned reads using featureCounts (v2.0.3) (83). Normalization and differential expression analysis, were performed using DESeq2 (84) (1.42.1). Unless otherwise stated, all reported p values have been adjusted for multiple testing using the Benjamini Hochberg procedure.

### Repeat element expression analysis

The analysis of repeat element expression from RNA-seq data began with the acquisition of raw sequencing reads in FASTQ format. Quality control was performed using FastQC (v0.11.9) to assess read quality, adapter contamination, and potential biases. Low-quality bases (Phred score < 20) and adapter sequences were trimmed using FASTP (0.32.2) (85).

Trimmed reads were then aligned to the human reference genome (hg38) using Bowtie2 (v2.5.1) with parameters optimized for sensitive mapping of repetitive elements (allowing multimapping reads). The alignment files (BAM) were subsequently processed using RepEnrich2, a pipeline designed for repeat element quantification. RepEnrich2 (86) partitions multimapping reads among repeat families using an expectation-maximization (EM) algorithm, enabling accurate family level expression estimation.

The resulting count tables of repeat element families were then imported into DESeq2 (1.42.1) for differential expression analysis. Normalization was performed using TMM (edgeR v4.0.16) to account for library size differences, and statistical testing identified significantly differentially expressed repeat elements (FDR < 0.05, |log2FC| > 1)

Functional interpretation involved classifying repeats by family (LINE, SINE, LTR, DNA transposons) and assessing their potential regulatory roles through coexpression analysis with neighboring genes (±10 kb) using Spearman correlation. Visualization of heatmaps and bar plots was done using ggplot2 (3.5.2).

### Clustering and visualization

Global changes in expression levels were evaluated by hierarchical clustering of samples and PCA using normalized expression data coupled with variance stabilized transformation. For hierarchical clustering, Euclidean distance was used as the distance metric. For visualization, normalized Bigwig tracks were generated using BEDtools (87) (v2.30.3). Integrative Genomic Viewer (88) was used for data visualization. Gene Set Enrichment Analysis (GSEA) was carried out with the GSEA (v 4.4.0) (89) to evaluate whether curated gene sets exhibit coordinated, statistically significant shifts between the two experimental conditions. For every gene set, an enrichment score (ES) was computed and then normalized (NES) to correct for gene set size. Statistical significance was determined using the recommended false discovery rate (FDR) cutoff of < 0.25. All enrichment plots and summary outputs were generated directly within the GSEA interface.

### Analysis of ChIP-seq data

#### Data processing, peak calling and annotations

Sequence reads were aligned to the human genome (hg38) using Bowtie2 (2.4.4) (90) under default settings. PCR duplicates were removed using PICARD tools generating, BAM files Read processing and alignment for analysis of repeat elements was performed separately (see below for details). Significant peaks were identified using Model Based Analysis for ChIP-seq (MACS3 3.0.2) (91) with a p value cutoff of 1e^−9^. Peaks were annotated using HOMER (v5.1) (92) with promoter regions classified as any peak within +/− 3 kb of a transcriptional start site (TSS), and enhancer region greater than 2.5 kb from a TSS. For visualization of H3K27ac profiles, BAM alignment files were normalized to RPKM values using DeepTools (3.5.1) (93) and visualized in Integrated Genome Viewer.

To investigate differential H3K27ac enrichment between L1 EN^+^ p53^-/-^ and L1 EN^-^ p53^-/-^ samples, we first identified unique and shared peaks using the narrowPeak files derived from peak calling. Peaks specific to L1 EN^+^ p53^-/-^ (gained peaks) were defined as those present in L1 EN^+^ p53^-/-^ but absent in L1 EN^-^ p53^-/-^, while peaks exclusive to L1 EN^-^ p53^-/-^ (lost peaks) were similarly defined. Additionally, we categorized a subset of the gained peaks as "newly gained" if they were located more than 5 kb away from any peak in the L1 EN^-^ p53^-/-^ sample, suggesting potential *de novo* enhancer activation. Peaks common to both samples were classified as maintained peaks. To quantify signal intensities over these peak categories, we extracted normalized ChIP-seq signal values from the corresponding bigWig files using deepTools. The average signal intensity for each region was computed for both samples. Subsequently, fold changes in signal between L1 EN^+^ p53^-/-^ and L1 EN^-^ p53^-/-^ were calculated for each peak, enabling comparative analysis of H3K27ac enrichment. Finally, these values were aggregated by category (gained, new gained, lost, maintained) to assess global shifts in acetylation and to visualize chromatin changes associated with each condition.

### Visualization of samples based on H3K27ac

Read coverage tracks were generated using deepTools (3.5.1) (bamCoverage with RPKM normalization) and visualized in the Integrative Genomics Viewer (IGV). Heatmaps and average profile plots were generated using computeMatrix and plotHeatmap to compare H3K27ac signals across samples.

### Differential H3K27ac enrichment analysis

Utilizing the read depth normalized matrix of H3K27ac signal for all consensus H3K27ac peaks, differential H3K27ac loci between A375 WT cells expressing L1 EN^+^ and p53^-/-^ cells expressing L1 EN^+^ and L1 EN^-^ was determined using DiffBind, employing the DESeq2 method. Significant regions were further filtered to events with an FDR < 0.05.

### Integration of ChIP-seq and RNA-seq Data

To investigate the regulatory relationship between H3K27ac marked regions and gene expression, we integrated ChIP-seq and RNA-seq data by linking differential H3K27ac peaks with nearby (±50 kb) differentially expressed genes. Spearman correlation analysis was performed to assess the association between H3K27ac signal intensity and gene expression levels. Additionally, we examined whether repeat elements with H3K27ac enrichment showed corresponding transcriptional changes in RNA-seq data, suggesting potential regulatory roles. Functional enrichment analysis was applied to H3K27ac associated repeats to identify relevant biological pathways.

### Data Availability

RNA-seq and ChIP-seq datasets acquired and described in our study have been deposited into the NCBI public database and are available under GEO Submission PRJNA1293546.

For analyses involving A375 WT and p53^-/-^ cells, the raw sequencing data were obtained from the publicly available GEO dataset GSE159134.

## Author information

### Corresponding author

Correspondence to Dr. Bhavana Tiwari.

## Acknowledgements

We sincerely thank Prof. Kathelen Burns (Harvard Medical School) for generously providing the pMT646 and pMT1093 plasmids used in this study. We are grateful to Mamuni Swain (iBRIC-ILS, Bhubaneswar) for her guidance in library preparation, and to Dr. Pramod Kumar Yadav (IISER Berhampur) for methodological support. We also acknowledge the Central Instrumentation Facility and the Department of Biological Sciences Instrumentation Facility at IISER Berhampur. Graphs were generated using GraphPad Prism, and illustrations were created with BioRender.

