## Supplemental Figure for "LINE-1 transposon derived DNA lesions reshape the chromatin landscape to promote genome instability"

**Running title:** LINE-1 ORF2p driven DNA damage and chromatin responses

Author name: Arun Kumar[1*], Ankita Subhadarsani Parida[1*], Kiran Kiran[1], Sunil K. Raghav[2], and Bhavana Tiwari[1#]

*Equal contribution

**Affiliations of all authors:**

[1]Department of Biological Sciences, Indian Institute of Science Education and Research (IISER) Berhampur, Odisha, India

[2]Immunogenomics & Systems Biology group, BRIC-Institute of Life Sciences (BRIC-ILS), Bhubaneswar, Odisha, India

**Keywords:** LINE-1 retrotransposon, ORF2p endonuclease, Double strand break, Genomic instability, Non homologous end joining (NHEJ), Enhancer remodelling.

**Figure EV1:**

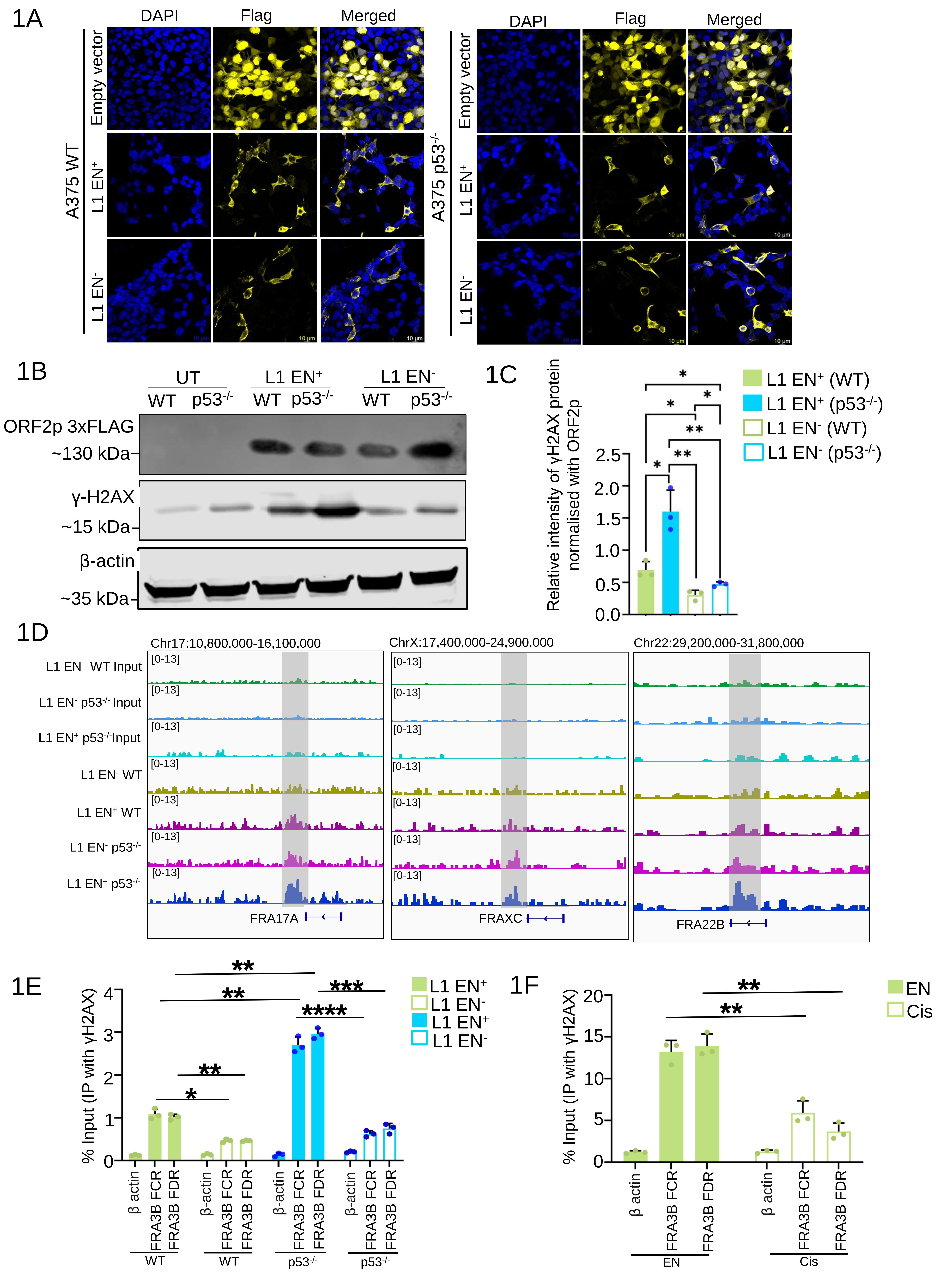
1A
DAPI
Flag
Merged
A375 WT
Empty vector
L1 EN+
L1 EN-
A375 p53-/-
10 µm
1B
UT
L1 EN+
L1 EN-
WT p53-/-
ORF2p 3xFLAG
~130 kDa
γ-H2AX
~15 kDa
β-actin
~35 kDa
1C
Relative intensity of γH2AX protein normalised with ORF2p
L1 EN+ (WT)
L1 EN+ (p53-/-)
L1 EN- (WT)
L1 EN- (p53-/-)
1D
Chr17:10,800,000-16,100,000
ChrX:17,400,000-24,900,000
Chr22:29,200,000-31,800,000
L1 EN+ WT Input
L1 EN- p53-/- Input
L1 EN+ p53-/-Input
L1 EN- WT
L1 EN+ WT
L1 EN- p53-/-
L1 EN+ p53-/-
[0-13]
FRA17A
FRAXC
FRA22B
1E
% Input (IP with γH2AX)
L1 EN+
L1 EN-
β actin
FRA3B FCR
FRA3B FDR
WT
p53-/-
1F
% Input (IP with γH2AX)
EN
Cis

**Figure EV1: Analysis of DNA damage induced by EN domain of L1-ORF2p**

**(1A)** Immunofluorescence microscopy was performed in A375 WT and $p53^{-/-}$ cells transfected with L1 $EN^{+}$ (pMT646), L1 $EN^{-}$ (pMT1093), or empty vector. Expression was detected using anti-FLAG antibody (Green), and nuclei were counterstained with DAPI (blue). Both L1 $EN^{+}$ and L1 $EN^{-}$ variants, as well as the empty vector, showed detectable expression. Images shown are representative of three independent biological replicates. Scale bars: 20 μm.

**(1B)** Western blotting was performed in A375 WT and $p53^{-/-}$ cells transfected with L1 $EN^{+}$ and L1 $EN^{-}$ constructs, along with non transfected controls. ORF2p expression was detected using anti FLAG antibody. γH2AX levels were used as a marker of DNA damage, while β-actin served as a loading control.

**(1C)** Densitometry quantification of γH2AX band intensities from immunoblots of A375 WT and $p53^{-/-}$ cells expressing L1 $EN^{+}$ and L1 $EN^{-}$ constructs. Band intensities were normalized to ORF2p. Quantification using Fiji represents mean ± SD from three independent biological replicates. Statistical analysis was performed using the Mann-Whitney U test, with significance indicated as *p < 0.05, **p < 0.01, and ns = not significant.

**(1D)** IGV snapshots of Fragile hotspot sites loci across L1 $EN^{-}$ and L1 $EN^{+}$ conditions in wild type and $p53^{-/-}$ A375 cells. Tracks display γ-H2AX ChIP seq signals. Shaded areas mark loci where elevated γ-H2AX signal aligns with increased transcript abundance.

**(1E)** Chromatin immunoprecipitation (ChIP) followed by qPCR was performed to assess the enrichment of γH2AX at fragile sites FRA3B of FCR and FDR region in U2OS cells transfected with either the active L1 endonuclease construct pMT646 ($EN^{+}$) or the catalytically inactive control pMT1093 ($EN^{-}$). The β actin locus was included as a negative control. Data are presented as mean ± standard deviation (S.D) from three independent biological replicates. Statistical analysis was conducted using the Mann Whitney U test; significance is indicated as p* < 0.05, p**<0.01, p***<0.001 and ns denotes non-significant comparisons.

**(1F)** Chromatin immunoprecipitation (ChIP) followed by qPCR was performed to evaluate γH2AX enrichment at the fragile site FRA3B (FCR and FDR regions) in A375 cells transfected with LINE1 endonuclease (EN) or treated with cisplatin. The β-actin locus was used as a negative control. Data are presented as mean ± standard deviation (SD) from three independent biological replicates. Statistical significance was assessed using the Mann Whitney U test, with p* < 0.05, p**<0.01, p***<0.001, and ns indicating non-significant differences.

**Figure EV2:**

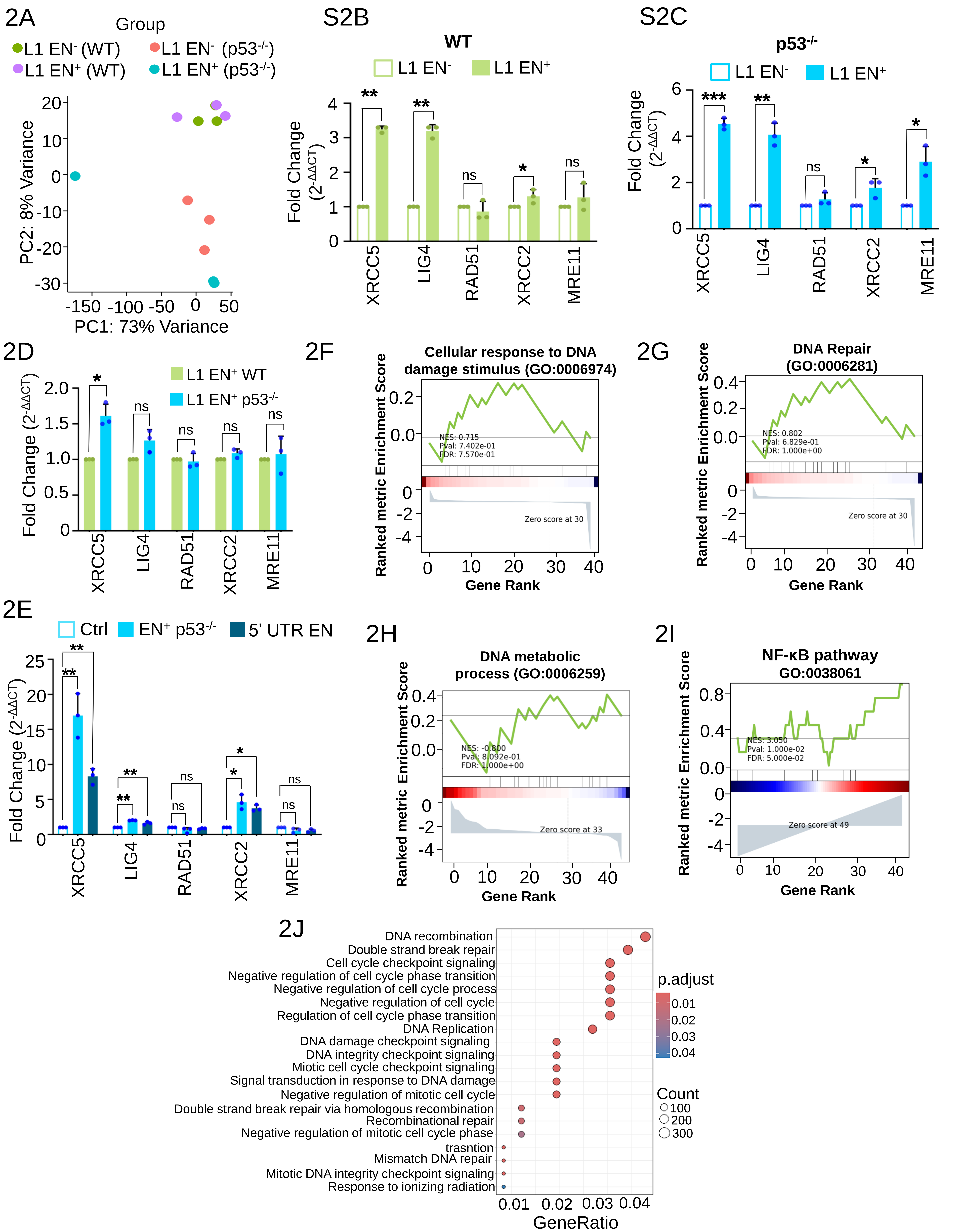

2A
Group
L1 EN- (WT)
L1 EN+ (WT)
L1 EN- (p53-/-)
L1 EN+ (p53-/-)
PC2: 8% Variance
PC1: 73% Variance
S2B
WT
L1 EN-
L1 EN+
Fold Change (2-ΔΔCT)
XRCC5
LIG4
RAD51
XRCC2
MRE11
S2C
p53-/-
2D
L1 EN+ WT
L1 EN+ p53-/-
2E
Ctrl
EN+ p53-/-
5' UTR EN
2F
Cellular response to DNA damage stimulus (GO:0006974)
NES: 0.715
Pval: 7.402e-01
FDR: 7.570e-01
Zero score at 30
Ranked metric Enrichment Score
Gene Rank
2G
DNA Repair (GO:0006281)
NES: 0.802
Pval: 6.829e-01
FDR: 1.000e+00
Zero score at 30
2H
DNA metabolic process (GO:0006259)
NES: -0.800
Pval: 8.092e-01
FDR: 1.000e+00
Zero score at 33
2I
NF-κB pathway GO:0038061
NES: 3.050
Pval: 1.000e-02
FDR: 5.000e-02
Zero score at 49
2J
DNA recombination
Double strand break repair
Cell cycle checkpoint signaling
Negative regulation of cell cycle phase transition
Negative regulation of cell cycle process
Negative regulation of cell cycle
Regulation of cell cycle phase transition
DNA Replication
DNA damage checkpoint signaling
DNA integrity checkpoint signaling
Miotic cell cycle checkpoint signaling
Signal transduction in response to DNA damage
Negative regulation of mitotic cell cycle
Double strand break repair via homologous recombination
Recombinational repair
Negative regulation of mitotic cell cycle phase trasntion
Mismatch DNA repair
Mitotic DNA integrity checkpoint signaling
Response to ionizing radiation
GeneRatio
p.adjust
Count

**Figure EV2: RNA seq comparison of EN deficient vs. EN proficient L1-ORF2p expression in WT and $p53^{-/-}$ melanoma Cells**

**(2A)** PCA plot illustrating clustering of global transcriptomic profiles across four experimental conditions: A375 WT and $p53^{-/-}$ cells transfected with either the catalytically inactive pMT1093 (L1 $EN^{-}$) or active pMT646 (L1 $EN^{+}$) constructs. Each point represents one of three independent biological replicates, with clear separation observed between groups, indicating condition specific transcriptional reprogramming associated with L1 EN activity and p53 status. One biological replicate from the $p53^{-/-}$ $EN^{+}$ group exhibited substantial variance relative to the other replicates and was therefore excluded from downstream analyses to maintain statistical rigor and data consistency.

**(2B,2C)** Bar plots displaying mRNA expression levels by qRT-PCR for XRCC5, LIG4, RAD51, XRCC2, and MRE11 in WT (D) and $p53^{-/-}$ (E) A375 cell lines expressing L1 $EN^{+}$ (filled) or L1 $EN^{-}$ (unfilled). The individual data points indicate three biological replicates, mean ± S.D., statistical significance was assessed using the Mann Whitney U test. Significance levels are indicated as follows: $*p < 0.05$, $**p < 0.01$, $***p < 0.001$, and ns = not significant.

**(2D)** Bar plots displaying the average qRT PCR expression levels (mean ± S.D.) for XRCC5, LIG4, RAD51, XRCC2, and MRE11 across three independent biological replicates in WT and $p53^{-/-}$ A375 cell lines expressing $L1EN^{+}$. Statistical significance was assessed using the Mann Whitney U test. Significance levels are indicated as follows: $*p < 0.05$, $**p < 0.01$, $***p < 0.001$, and ns = not significant.

**(2E)** Bar plots displaying mRNA expression levels by qRT-PCR for for XRCC5, LIG4, RAD51, XRCC2, PARP1 and MRE11 in $p53^{-/-}$ A375 cell lines expressing empty vector control, APE1 and EN-GFP. The individual data points indicate three biological replicates, mean ± S.D., statistical significance was assessed using the Mann Whitney U test. Significance levels are indicated as follows: $*p < 0.05$, $**p < 0.01$, $***p < 0.001$, and ns = not significant.

**(2F-2I):GO:0006974 (cellular response to DNA damage stimulus)**: Negatively enriched in WT $L1EN^{-}$.

• **GO:0006259 (DNA metabolic process)**: Not significant in WT $L1EN^{-}$ .

• **GO:0006281 (DNA repair):** Downregulated in WT $L1EN^{-}$.

• **GO:0038061 (NF-κB pathway):** Significantly upregulated in WT L1 EN+ (NES = 3.05, P = $1.00 \times 10^{-2}$, FDR = $5.00 \times 10^{-2}$).Each plot shows the enrichment score curve (green), gene rank positions (black lines), and the distribution of enrichment signals (bottom heatmap).

**(2J)** A dot summarizing enriched pathways related to DNA repair genes. The x axis represents the GeneRatio (proportion of genes in the pathway), and the y axis lists the pathways (e.g., DNA recombination, double strand break repair). Dot size or bar height indicate the count of genes in each pathway, and color intensity reflects the adjusted p value (e.g., 2.5e-12, 5.0e-12).

### Figure EV3:

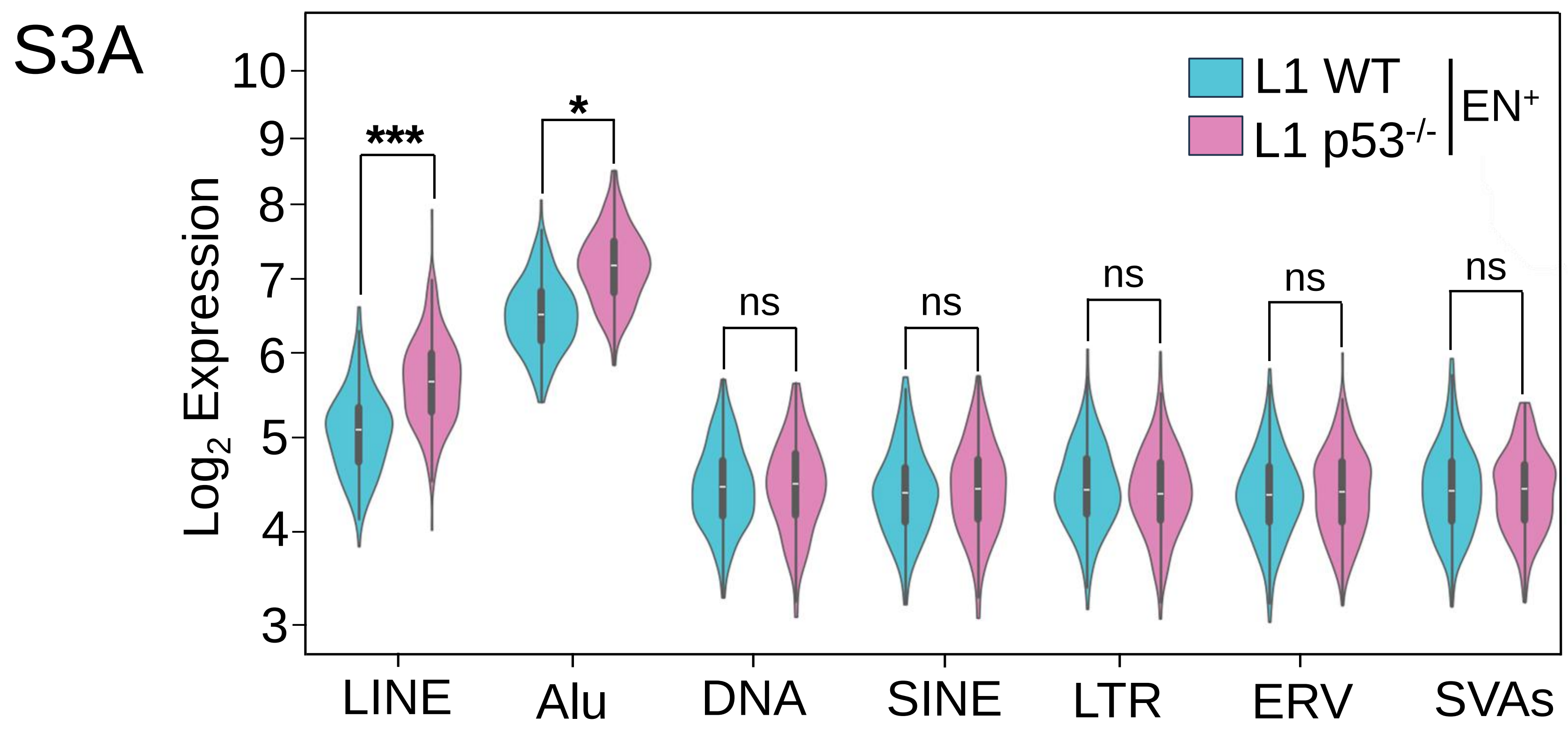


**Figure EV3: Transcriptomic profiling of transposable element families in A375 wild type and p53$^{-/-}$ cells expressing L1 EN$^{+}$**

**(3A)** Violin plot showing the expression distribution of selected transposable element families in A375 WT cells expressing L1EN$^{+}$ and p53$^{-/-}$ expressing L1 EN$^{+}$ p53$^{-/-}$ samples. Each violin represents the distribution across biological replicates, with individual dots indicating replicate level values and internal box plots denoting the median and interquartile range. Statistical significance was assessed using an unpaired two tailed t test; with "ns" denoting non significant differences ($p > 0.05$).

**Figure EV4:**

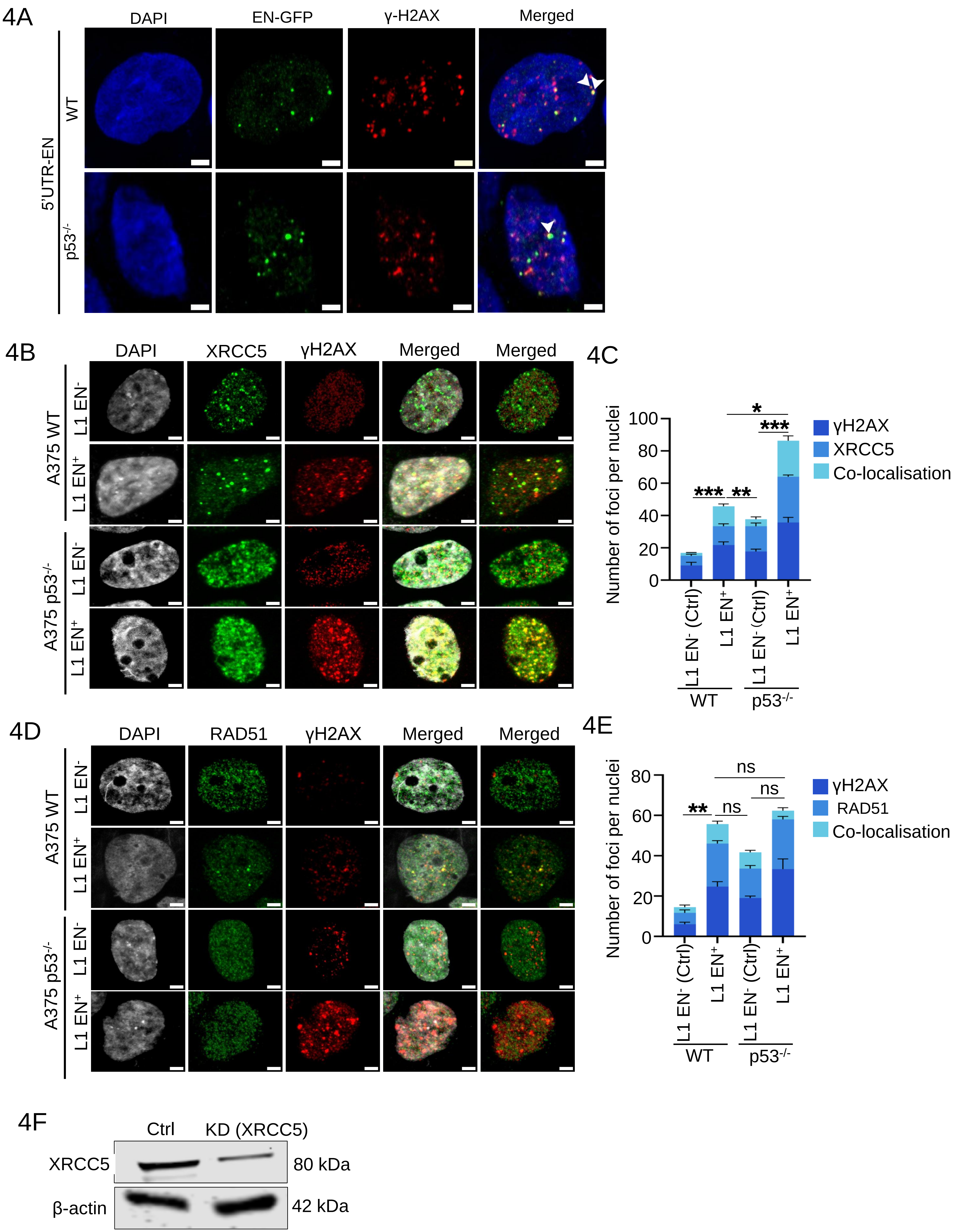
4A
DAPI
EN-GFP
γ-H2AX
Merged
5'UTR-EN
WT
p53-/-
4B
DAPI
XRCC5
γH2AX
Merged
Merged
A375 WT
L1 EN-
L1 EN+
A375 p53-/-
L1 EN-
L1 EN+
4C
Number of foci per nuclei
100
80
60
40
20
0
*
***
***
**
γH2AX
XRCC5
Co-localisation
L1 EN- (Ctrl)
L1 EN+
L1 EN- (Ctrl)
L1 EN+
WT
p53-/-
4D
DAPI
RAD51
γH2AX
Merged
Merged
A375 WT
L1 EN-
L1 EN+
A375 p53-/-
L1 EN-
L1 EN+
4E
Number of foci per nuclei
80
60
40
20
0
ns
ns
**
ns
γH2AX
RAD51
Co-localisation
L1 EN- (Ctrl)
L1 EN+
L1 EN- (Ctrl)
L1 EN+
WT
p53-/-
4F
Ctrl
KD (XRCC5)
XRCC5
80 kDa
β-actin
42 kDa

**Figure EV4: Colocalisation analysis of γH2AX with XRCC5 and RAD51 in LI EN+ and L1 EN- in A375 WT and $p53^{-/-}$ cells**

**(4A)** Representative confocal images of A375 WT and $p53^{-/-}$ cells transfected with GFP tagged endonuclease only domain (EN-GFP/EN, green) expressed under the control of L1-5'UTR promoter, immunostained for the DNA damage marker γH2AX (red), and counterstained with DAPI (blue) to visualize nuclei (A). Immunofluorescence was performed to assess the spatial co-localization of EN expression and DNA damage foci. EN localization was detected using an anti-GFP antibody, while γH2AX was visualized using an anti phospho H2AX (Ser139) antibody. Scale bar: 2 µm.

**(4B)** Immunofluorescence staining of XRCC5 (green) and γH2AX (red) in A375 WT and $p53^{-/-}$ cells transfected with L1 $EN^{+}$ or L1 $EN^{-}$. Nuclei are counterstained with DAPI (blue). Scale bar: 2 µm.

**(4D)** Immunofluorescence staining of γH2AX (red) and RAD51 (green) in A375 WT and $p53^{-/-}$ cells expressing L1 $EN^{+}$ or L1 $EN^{-}$. DAPI (blue) stains nuclei. Scale bar: 2 µm.

**(4C, 4E)** Quantification of γH2AX with XRCC5 (H) , and γH2AX with RAD51(J) foci and their colocalized signals in A375 WT and $p53^{-/-}$ cells transfected with L1 $EN^{+}$ or L1 $EN^{-}$. Data are shown as stacked bar plots representing mean foci counts from three biological replicates (30 fields per replicate). Statistical significance was assessed using the Mann Whitney U test, colocalization was determined by Pearson correlation coefficient (r = 0.6–1). The data represents mean ± SD across three biological replicates. Statistical significance was evaluated using the Mann Whitney U test, with significance levels represented as follows: *p<0.05, **p<0.01 and ***p<0.001.

**(4F)** Representative immunoblot analysis of A375 cells transiently transfected with TLV2 vector control or TLV2-CRISPR plasmid targeting XRCC5. Whole-cell lysates were probed with an anti-XRCC5 antibody to assess knockdown efficiency. β-Actin was used as a loading control.

**Figure EV5:**

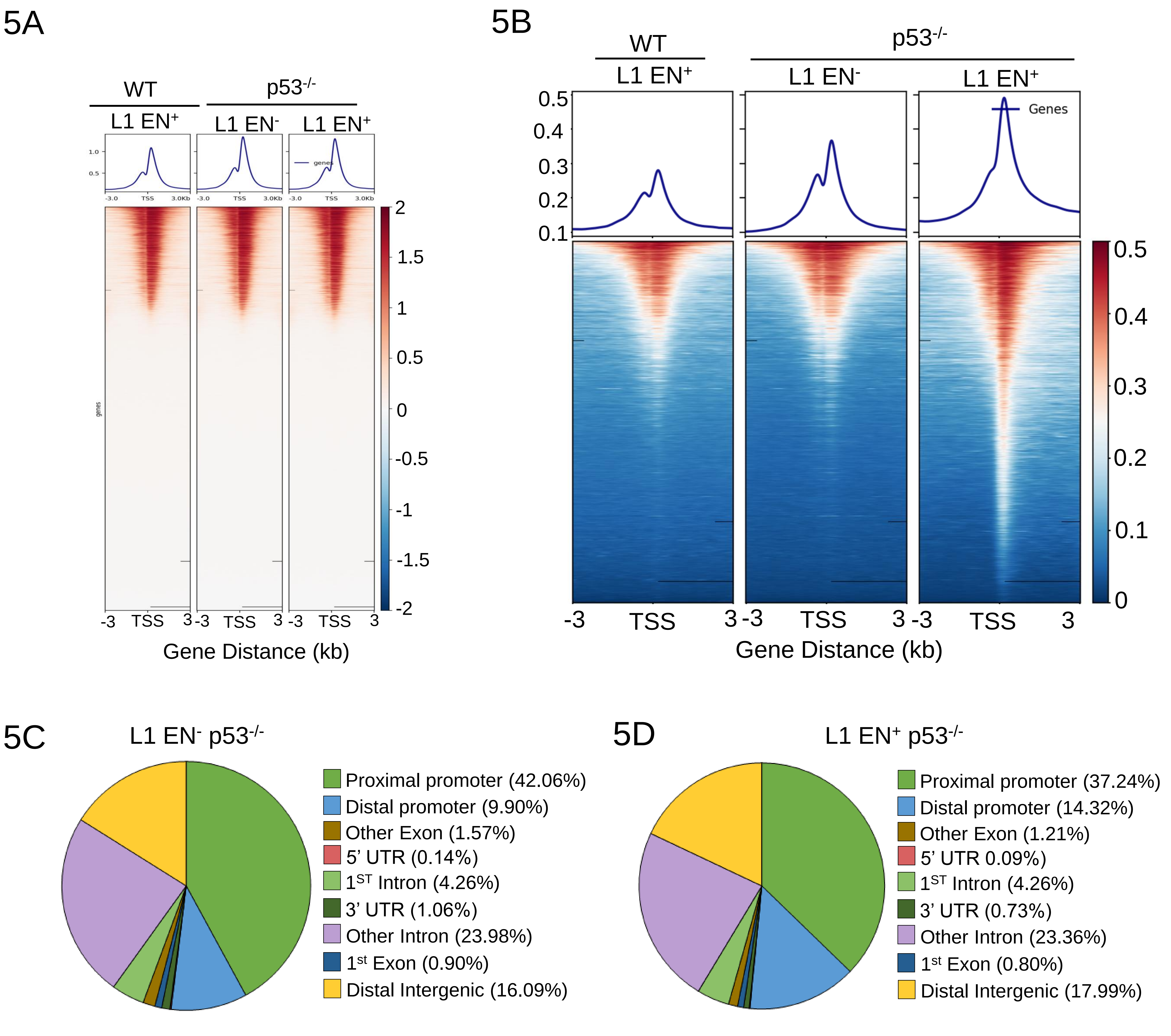


**Figure EV5: H3K4me3 and H3K27ac chIP-seq analyses in L1 EN$^+$ vs EN$^-$ A375 p53$^{-/-}$ cells**

**(5A)** This heatmap displays the normalized H3K4me3 ChIP seq signal intensity ±3 kb around the TSS of annotated genes using data from A375 WT expressing L1 EN$^+$ WT and A375 p53$^{-/-}$ cells expressing L1 EN$^-$ and L1 EN$^+$ p53$^{-/-}$. Red indicates higher enrichment and blue indicates lower enrichment.

**(5B)** This heatmap displays the normalized H3K27ac ChIP seq signal intensity ±3 kb around the TSS of annotated genes using data from A375 WT expressing L1 EN$^+$ WT, and p53$^{-/-}$ cells expressing L1 EN$^-$ and L1 EN$^+$ p53$^{-/-}$. The signal matrix was generated using computeMatrix in reference point mode and visualized with a diverging color map ("RdBu_r"), where red indicates higher enrichment and blue indicates lower enrichment.

**(5C and D)** Pie charts depicting the genomic distribution of H3K27ac enriched regions in A375 p53$^{-/-}$ cells expressing (A) L1 EN$^-$ and (B) L1 EN$^+$. H3K27ac marked loci indicative of active promoters and enhancers were annotated to genomic features including promoters, 5′UTRs, 3′UTRs, first exons, other exons, first introns, other introns, and distal intergenic regions based on proximity to RefSeq genes. Color coded segments denote feature-specific annotations as indicated in the accompanying legend.

**Figure EV6:**

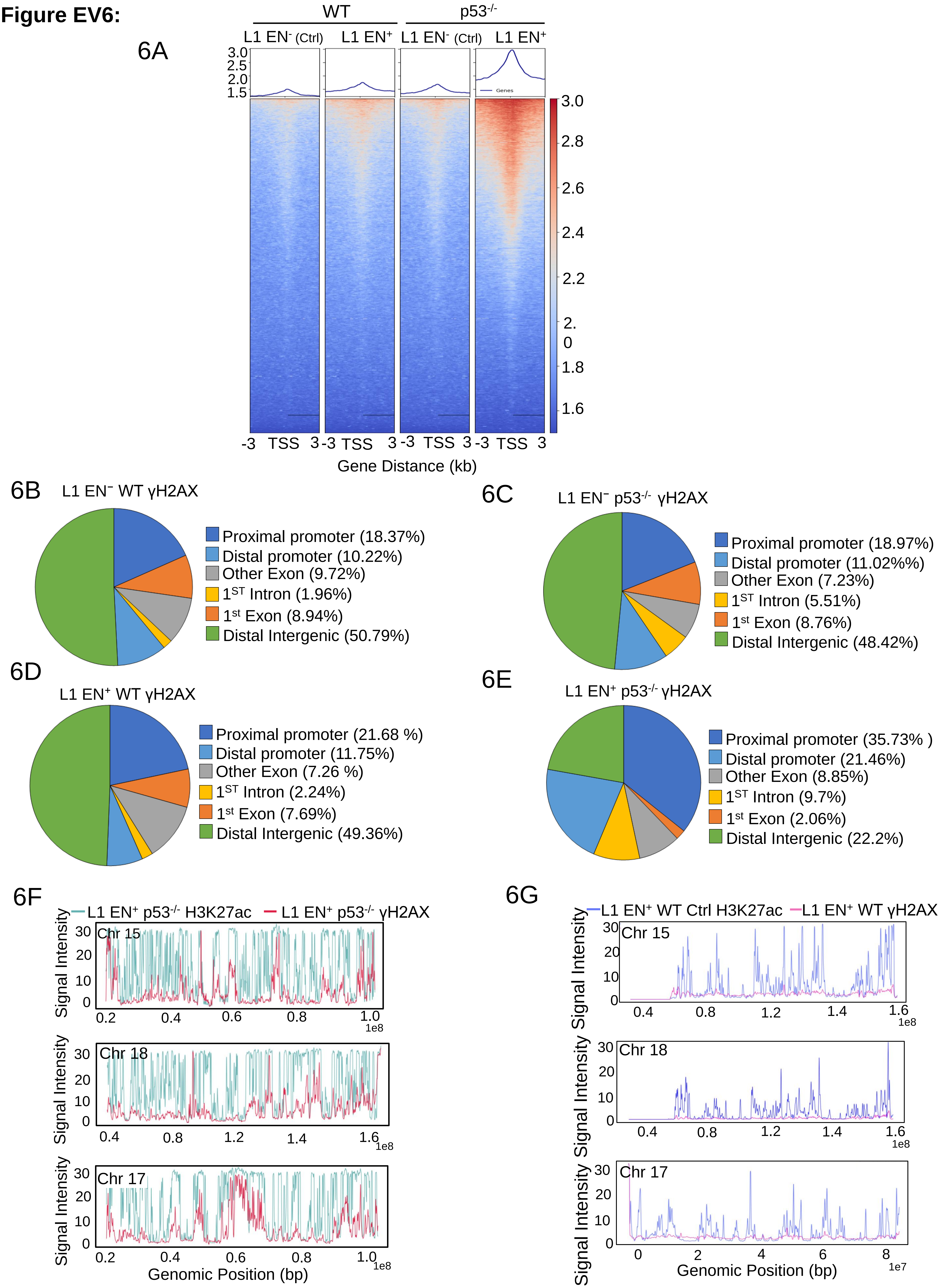

6A
WT
p53-/-
L1 EN- (Ctrl)
L1 EN+
L1 EN- (Ctrl)
L1 EN+
Genes
-3 TSS 3
Gene Distance (kb)
6B
L1 EN− WT γH2AX
Proximal promoter (18.37%)
Distal promoter (10.22%)
Other Exon (9.72%)
1ST Intron (1.96%)
1st Exon (8.94%)
Distal Intergenic (50.79%)
6C
L1 EN− p53-/- γH2AX
Proximal promoter (18.97%)
Distal promoter (11.02%%)
Other Exon (7.23%)
1ST Intron (5.51%)
1st Exon (8.76%)
Distal Intergenic (48.42%)
6D
L1 EN+ WT γH2AX
Proximal promoter (21.68 %)
Distal promoter (11.75%)
Other Exon (7.26 %)
1ST Intron (2.24%)
1st Exon (7.69%)
Distal Intergenic (49.36%)
6E
L1 EN+ p53-/- γH2AX
Proximal promoter (35.73% )
Distal promoter (21.46%)
Other Exon (8.85%)
1ST Intron (9.7%)
1st Exon (2.06%)
Distal Intergenic (22.2%)
6F
L1 EN+ p53-/- H3K27ac
L1 EN+ p53-/- γH2AX
Signal Intensity
Chr 15
Chr 18
Chr 17
Genomic Position (bp)
1e8
6G
L1 EN+ WT Ctrl H3K27ac
L1 EN+ WT γH2AX
Signal Intensity
Chr 15
Chr 18
Chr 17
Genomic Position (bp)
1e8
1e7

**Figure EV6: Overlaying H3K27ac and γH2AX datasets**

(**6A**) Heatmap showing normalized γH2AX ChIP-seq signal intensity ±3 kb around the transcription start sites (TSS) of annotated genes in A375 WT and $p53^{-}/^{-}$ cells expressing either L1 $EN^{-}$ and L1 $EN^{+}$. Red indicates increased enrichment and blue indicates reduced enrichment.

**(6B and E)** Pie charts depicting the genomic distribution of H3K27ac enriched regions in A375 $p53^{-/-}$ cells expressing (A) L1 $EN^{-}$ and (B) L1 $EN^{+}$. H3K27ac marked loci indicative of active promoters and enhancers were annotated to genomic features including promoters, 5′UTRs, 3′UTRs, first exons, other exons, first introns, other introns, and distal intergenic regions based on proximity to RefSeq genes. Color coded segments denote feature-specific annotations as indicated in the accompanying legend.

**(6F)** Genome wide distribution of ChIP-seq signal intensities for H3K27ac (blue) and γH2AX (red) along chromosomes 15, 18 and 17 in L1 $EN^{+}$ WT A375 cells. Signal values exceeding 30 bins were compressed to enhance visualization of overall peak distribution. The x-axis denotes genomic position (bp), and the y-axis represents normalized signal intensity.

**(6G)** Genome-wide distribution of ChIP-seq signal intensities for H3K27ac (blue) and γH2AX (red) along chromosomes 15, 18 and 17 in L1 $EN^{+}$ $p53^{-/-}$ A375 cells. Signal values exceeding 30 bins were compressed to improve visualization. The x-axis denotes genomic position (bp), and the y-axis represents normalized signal intensity.

**Figure EV7:**

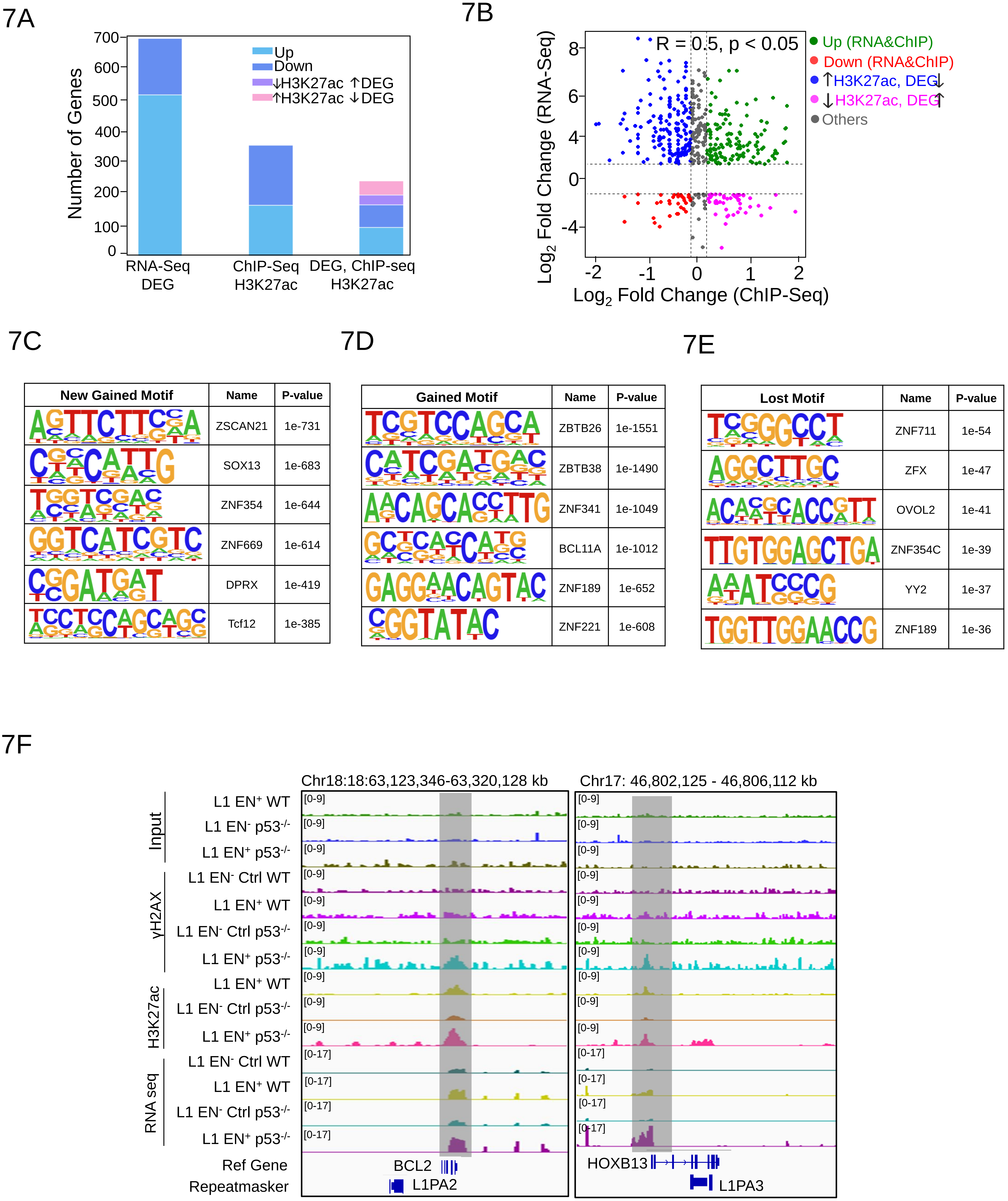
7A
Number of Genes
700
600
500
400
300
200
100
0
Up
Down
↓H3K27ac ↑DEG
↑H3K27ac ↓DEG
RNA-Seq DEG
ChIP-Seq H3K27ac
DEG, ChIP-seq H3K27ac
7B
R = 0.5, p < 0.05
Up (RNA&ChIP)
Down (RNA&ChIP)
↑H3K27ac, DEG↓
↓H3K27ac, DEG↑
Others
Log2 Fold Change (RNA-Seq)
Log2 Fold Change (ChIP-Seq)
7C
New Gained Motif
Name
P-value
ZSCAN21 1e-731
SOX13 1e-683
ZNF354 1e-644
ZNF669 1e-614
DPRX 1e-419
Tcf12 1e-385
7D
Gained Motif
Name
P-value
ZBTB26 1e-1551
ZBTB38 1e-1490
ZNF341 1e-1049
BCL11A 1e-1012
ZNF189 1e-652
ZNF221 1e-608
7E
Lost Motif
Name
P-value
ZNF711 1e-54
ZFX 1e-47
OVOL2 1e-41
ZNF354C 1e-39
YY2 1e-37
ZNF189 1e-36
7F
Chr18:18:63,123,346-63,320,128 kb
Chr17: 46,802,125 - 46,806,112 kb
Input
γH2AX
H3K27ac
RNA seq
L1 EN+ WT
L1 EN- p53-/-
L1 EN+ p53-/-
L1 EN- Ctrl WT
L1 EN+ WT
L1 EN- Ctrl p53-/-
L1 EN+ p53-/-
L1 EN+ WT
L1 EN- Ctrl p53-/-
L1 EN+ p53-/-
L1 EN- Ctrl WT
L1 EN+ WT
L1 EN- Ctrl p53-/-
L1 EN+ p53-/-
Ref Gene
Repeatmasker
BCL2
L1PA2
HOXB13
L1PA3

**Figure EV7:Integrative analysis of RNA seq and H3K27ac ChIP seq**

**(7A)** Stacked bar plot classifying the RNA seq data into two categories: Upregulated (n = 560), Downregulated (n = 197), then the ChIP seq region into two categories, Upregulated (n = 179), Downregulated (n = 207) and the 115 common genes between RNAseq and H3K27ac ChIp seq datasets into four categories: Upregulated (RNA↑ & ChIP↑, n = 22), Downregulated (RNA↓ & ChIP↓, n = 14), Discordant (RNA↑ & ChIP↓, n = 19 and RNA↓ & ChIP↑, n = 12), and Others (n = 48).

**(7B)** Concordant Changes in gene expression and H3K27ac occupancy in L1 $EN^+$ WT vs L1 $EN^+$ $p53^{-/-}$ cells. This scatter plot illustrates the correlation between RNA seq ($log_2$ fold change in gene expression, y axis) and H3K27ac ChIP seq ($log_2$ fold change in H3K27ac signal, x axis) across individual genes. Genes are colored based on their regulation patterns: green indicates upregulation in both RNA seq and H3K27ac (UpBoth), red indicates downregulation in both (DownBoth), blue represents genes upregulated in RNA seq but downregulated in H3K27ac (UpRNA_DownChIP), magenta shows downregulation in RNA-seq but upregulation in H3K27ac (DownRNA_UpChIP), and gray marks non concordant or insignificant changes.

**(7C-E)** Sequence logos representing enriched DNA motifs identified in genomic regions showing differential H3K27ac modifications (New gained, gained and lost). Each row corresponds to a specific motif, with the height of each nucleotide reflecting its relative frequency at that position. The analysis was performed using a motif discovery tool, and all displayed motifs are significantly enriched with p-values< 1e-5, indicating strong statistical confidence in the motif overrepresentation compared to background regions.

**(7F)** IGV snapshots of BCL2 and HOXB13 loci across L1 $EN^-$ and L1 $EN^+$ conditions in wild-type and $p53^{-/-}$ A375 cells. Tracks display H3K27ac ChIP-seq and RNA-seq signals. Shaded areas of BCL2 and HOXB13 mark loci where elevated H3K27ac signal aligns with increased transcript abundance, supporting the hypothesis that L1-ORF2p endonuclease activity modulates enhancer driven gene expression.
